# Inference of Selective Force on House Mouse Genomes during Secondary Contact in East Asia

**DOI:** 10.1101/2023.08.07.552211

**Authors:** Kazumichi Fujiwara, Shunpei Kubo, Toshinori Endo, Toyoyuki Takada, Toshihiko Shiroishi, Hitoshi Suzuki, Naoki Osada

**Affiliations:** Mouse Genomics Resource Laboratory, National Institute of Genetics, Mishima, Japan; Graduate School of Information Science and Technology, Hokkaido University, Sapporo, Japan; Integrated BioResource Information Division, RIKEN BioResource Research Center, Tsukuba, Japan; RIKEN BioResource Research Center, Tsukuba, Japan; Graduate School of Environmental Science, Hokkaido University, Sapporo, Japan

## Abstract

The house mouse (*Mus musculus*), commensal to humans, has spread globally via human activities, leading to secondary contact between genetically divergent subspecies. This pattern of genetic admixture can provide insights into the selective forces at play in this well-studied model organism. Our analysis of 163 house mouse genomes, with a particular focus on East Asia, revealed substantial admixture between the subspecies *castaneus* and *musculus*, particularly in Japan and southern China. We revealed, despite the admixture, all Y chromosomes in the East Asian samples belonged to the *musculus*-type haplogroup, potentially explained by genomic conflict under sex ratio distortion due to varying copy numbers of ampliconic genes on sex chromosomes. We also investigated the influence of selection on the post-hybridization of the subspecies *castaneus* and *musculus* in Japan. Even though the genetic background of most Japanese samples closely resembles the subspecies *musculus*, certain genomic regions overrepresented the *castaneus*-like genetic components, particularly in immune-related genes. Furthermore, a large genomic block containing a vomeronasal/olfactory receptor gene cluster predominantly harbored *castaneus*-type haplotypes in the Japanese samples, highlighting the crucial role of olfaction-based recognition in shaping hybrid genomes.

## Introduction

The house mouse (*Mus musculus*) is a pivotal laboratory animal, which is commensal to humans and has spread worldwide due to human activities such as colonization, agricultural expansion, and trade (GÜndüz et al. 2001; Geraldes et al. 2008; Searle et al. 2009a; Searle et al. 2009b; Gabriel et al. 2011; Macholán et al. 2012; Jones et al. 2013; Li et al. 2020). The house mouse exhibits high morphological and genetic diversity, with three main subspecies, recognized: *M. musculus castaneus*, *M. m. musculus*, and *M. m. domesticus* (Boursot et al. 1996; Salcedo et al. 2007; Phifer-Rixey et al. 2020; Fujiwara et al. 2022a). These subspecies diverged around 180– 500 kya and underwent secondary contact after dispersing from their native habitat around South Asia (Boursot et al. 1993; Din et al. 1996) with asymmetric hybrid incompatibility; the subspecies *domesticus* shows relatively strong hybrid incompatibility with the other two subspecies (White et al. 2011; White et al. 2012). Despite partial reproductive isolation between subspecies, genetic studies have revealed a non-negligible amount of gene flow, and occasional identification of hybrid genotypes (Orth et al. 1998; Ďureje et al. 2012; Fujiwara et al. 2022a). Studying the genetic differentiation and hybridization patterns in *M. musculus* thus provides a unique opportunity to gain insights into human history and the genetic architecture of hybridization among genetically distinct subspecies of a well-studied model organism.

In East Asia, two major subspecies have been widely observed; the subspecies *castaneus* is distributed in southern China and Taiwan, and the subspecies *musculus* distributed in northern China, the Korean Peninsula, and the Russian Far East (Din et al. 1996; Jing et al. 2014; Fujiwara et al. 2022a). Another subspecies, *M. m. molossinus*, has been recognized in the Japanese archipelago, where researchers have shown through genetic analysis that it is derived from a hybrid between the subspecies *castaneus* and *musculus* (Yonekawa et al. 1980; Yonekawa et al. 1982; Yonekawa et al. 1988; Sakai et al. 2005; Takada et al. 2013), while most of wild Japanese house mice have major genetic background of the subspecies *musculus* (Fujiwara et al. 2022a).

Interestingly, mitochondrial haplogroups representing the subspecies *castaneus* are mainly distributed in northern Japan, while mitochondrial haplogroups representing the subspecies *musculus* are observed in southern Japan (Terashima et al. 2006; Kuwayama et al. 2017; Li et al. 2020), leading the hypothesis that the subspecies *castaneus* was introduced from southern China to the Japanese archipelago and *musculus* was introduced from the Korean Peninsula with wet-rice field cultivation system (Terashima et al. 2006; Kuwayama et al. 2017). This hypothesis would be similar to the well-known model of the origin of modern Japanese, called the dual structure model by Hanihara (1991). The dual structure model assumes that the genetic makeup of modern Japanese has been composed of both indigenous Jomon hunter-gatherers and newly migrated Yayoi rice farmers (Hanihara 1991). While recent genetic analyses of modern and ancient human genome studies have revealed that the Jomon originated from one of the basal East Asian populations (McColl et al. 2018; Kanzawa-Kiriyama et al. 2019; Gakuhari et al. 2020; Osada and Kawai 2021), it is largely unknown when and from where the subspecies *castaneus* arrived in the Japanese archipelago.

Although the genetic diversity of maternally-inherited mitochondrial haplotypes in wild house mice has been well studied, natural variation in the paternally-inherited Y chromosome was largely unknown except for a few examples (Morgan and Pardo-Manuel de Villena 2017). Recent experimental studies have discovered X-and Y-linked loci that are crucial for hybrid incompatibility, where copy number variation (CNV) plays a significant role in determining hybrid incompatibility and sex-ratio distortion (SD). Previous studies have demonstrated that Y-linked *Sly* suppresses post-meiotic expression of X-and Y-linked genes in sperms (Ellis et al. 2011). Copy number increase in *Sly* results in male-biased progenies, while copy number increase in X-linked *Slx1* are known to counteract it (Scavetta and Tautz 2010; Cocquet et al.). Morgan et al. (2017) showed that the subspecies *musculus* tended to have higher *Sly*/*Slx* copy numbers than other subspecies, but their analyzed samples are biased toward wild-derived inbred lines [31]. Therefore, the analysis for CNVs of *Sly*/*Slx* and their natural distribution range in wild samples would provide how the sex-ratio distortion alleles affect the genetic differentiation of mouse subspecies.

The selective forces on the formation of hybrid genomes are also an important issue in evolutionary biology research. With the advancement of genome sequencing technology, many genome-scale studies have been conducted to examine the effects of natural selection under interspecific or intersubspecific gene flow. In particular, adaptive gene introgression, in which beneficial alleles cross the boundary of populations and spread rapidly to another population, has attracted the interest of many researchers (Song et al. 2011; Pardo-Diaz et al. 2012; Huerta-Sánchez et al. 2014; Enard and Petrov 2018). The distinctive genetic characteristic of wild Japanese house mice would provide a great opportunity to investigate the effects of selection during hybridization. Having the major genetic background of the subspecies *musculus*, the preference for the genomic region of the *castaneus*-ancestry in the Japanese population leaves a similar signature to that of adaptive introgression and is a robust signal of natural selection during hybrid genome formation.

In this study, we analyze 163 high-coverage whole genome sequences of *M. musculus*, including newly sequenced samples primarily from Japan, to quantify the pattern of genetic admixture among the samples. By contrasting the population differentiation patterns of autosomal and sex-linked genomic regions, we infer the evolutionary history of the two East Asian subspecies populations. We also analyze Japanese samples to identify specific regions of the genome where genetic components from the subspecies *castaneus* are more prevalent, even though the majority of the Japanese samples have a genetic background from the subspecies *musculus*. Focusing on the evolutionary history of the *M. musculus* in East Asia, this study provides new insights into how the commensal animal has shaped its genetic traits through secondary contact mediated by human activities.

## Result

### Genome-wide pattern of differentiation in East Asia

In this study, we sequenced the genome of 37 wild house mice mainly collected from Japan, with an average coverage of 26.3 (Supplementary Table 1). Those data were merged with a previously published dataset that included *M. spretus*, yielding the genotypes 170 samples. After filtering, 133,886,237 autosomal biallelic SNVs were used for the initial population study.

Principal component analysis (PCA) was performed on all *M. musculus* samples (Supplementary Figure 1). The results were in high agreement with those previously shown by Fujiwara et al. (2022). Three major genetic components corresponding to subspecies *castaneus*, *domesticus*, and *musculus*, were observed. Most of the East Asian samples aligned on the *castaneus*-*musculus* cline, which means that the genetic characteristics of the East Asian samples can be modeled by different degrees of admixture between the subspecies *castaneus* and *musculus*. Genetic clustering was performed using ADMIXTURE software assuming three ancestral populations (*K* = 3). The results yielded three genetic components corresponding to the three main subspecies, consistent with the PCA results (Supplementary Figure 2).

To quantify the degree of genetic admixture in each sample, the *f*_4_-ratio value developed by Patterson et al. (2012) was calculated for each sample (see Methods), assuming that the East Asian samples were derived from admixture of two unknown lineages that shared ancestry with the Korean *musculus* and Indian *castaneus* populations. The estimated percentage of *musculus* ancestry, expressed as α, varied from 0.109 to 1 among the East Asian samples, as shown in Fig. 1. In the Japanese archipelago, all samples had α values greater than 0.564, and the mean value of α was 0.878 (Fig. 1). Interestingly, the samples from the Sea of Japan side have lower α values than those from the Pacific Ocean side. When the sample from Okinawa (a subtropical island) was excluded and the other samples were divided into two classes according to the Central Watershed of Japan, the Pacific-side samples had significantly higher α than the Sea of Japan-side samples (*P* = 0.0002, MWU test). In China, α values were close to 1 in northern and western China, but much lower in southern China (Fig. 1). In the southern Chinese samples, the averaged α was 0.295.

**Fig. 1:**
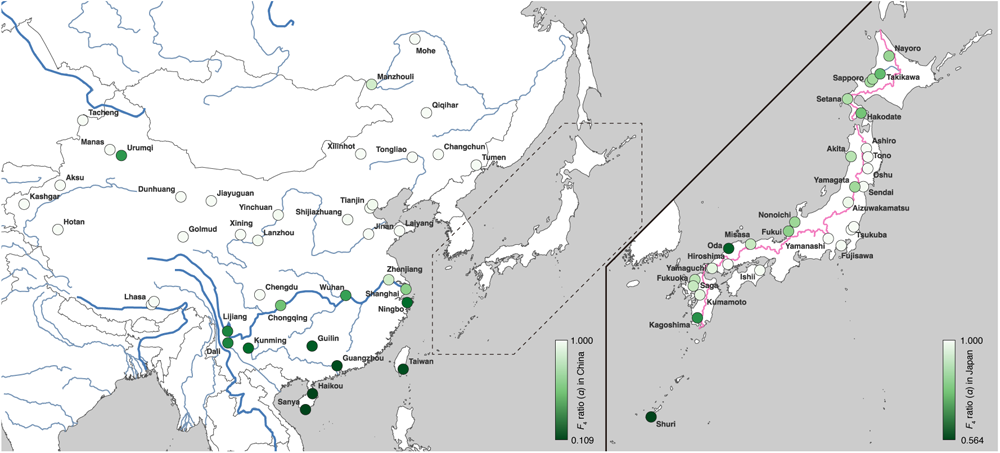
The estimated ratio of *musculus/castaneus* ancestry in East Asia.

The colors of circles indicate *musculus*/*castaneus* ancestry of samples at each collection site (left pane: East Asian samples except Japanese archipelago, right pane: Japanese archipelago samples). The *musculus* ancestry and *castaneus* ancestry of each sample are represented by a range of colors from white to green, with the stronger green color indicating higher *castaneus* ancestry. The range of ancestry ratio is indicated by the color bar in each panel. The pink line in Panel B represents the Central Watershed in the Japanese archipelago.

### Genetic differentiation at mitochondrial and sex-linked loci

We compared genetic differentiation patterns at autosomal, X-chromosomal, Y-chromosomal, and mitochondrial loci. We observed generally less mixing of subspecies genomes on the X chromosome; *i*.*e*., there was less *castaneus*-ancestry in the Japanese population and less *musculus*-ancestry in the southern Chinese population. The estimated α values on the X chromosome ranged from 0.317 to 1 (average α: 0.860) for Japanese samples and from 0.034 to 0.952 (average α: 0.207) for southern Chinese samples, showing that the X chromosome has a low level of introgression into the predominant genome. The difference in α between autosomes and X chromosomes were significant in both Japanese (*P* = 0.0008) and southern Chinese (*P* = 0.0134) populations (Wilcoxon signed rank test).

Genealogies of the Y-chromosomal and mitochondrial loci inferred by the maximum likelihood (ML) method are shown in Supplementary Figure 3 and 4. The Y-chromosomal locus formed three distinct haplogroups corresponding to the three major subspecies, whereas the mitochondrial locus showed five major haplogroups. In addition to the three mitochondrial haplogroups representing the subspecies *castaneus*, *domesticus,* and *musculus*, samples from Madagascar and Nepal formed distinct mitochondrial haplogroups, as previously shown (Fujiwara et al. 2022a; Fujiwara et al. 2022b).

The geographic distribution of the subspecies *castaneus* and *musculus* haplotypes was markedly different between mitochondrial and Y-chromosomal loci (Fig. 2). The distribution of mitochondrial haplotypes was consistent with previous findings (Fig. 2a). In the Japanese archipelago, *castaneus-* and *musculus-*type mitochondrial haplotypes are distributed in the northern and southern regions of mainland Japan, respectively. In China, the distribution of mitochondrial haplotypes was clearly divided around the Yangtze River basin (or divided at Qinling Mountains), with *musculus*-type haplotypes in northern China and *castaneus*-type haplotypes in southern China. However, all Y-chromosomal haplotypes in East Asia were assigned exclusively to *musculus*-type haplotypes (Fig. 2b). The Y chromosome of the reference genome (C57BL/6J) also had a *musculus*-type haplotype, and it was closely related to the haplotype of Japanese samples from the Tohoku area (Supplementary Figure 4), supporting the idea that the Y chromosome of classical inbred strains originated from Japanese house mice (Japanese fancy mouse) (Bishop et al. 1985; Nagamine et al. 1992; Tucker et al. 1992; Takada et al. 2013).

**Fig. 2:**
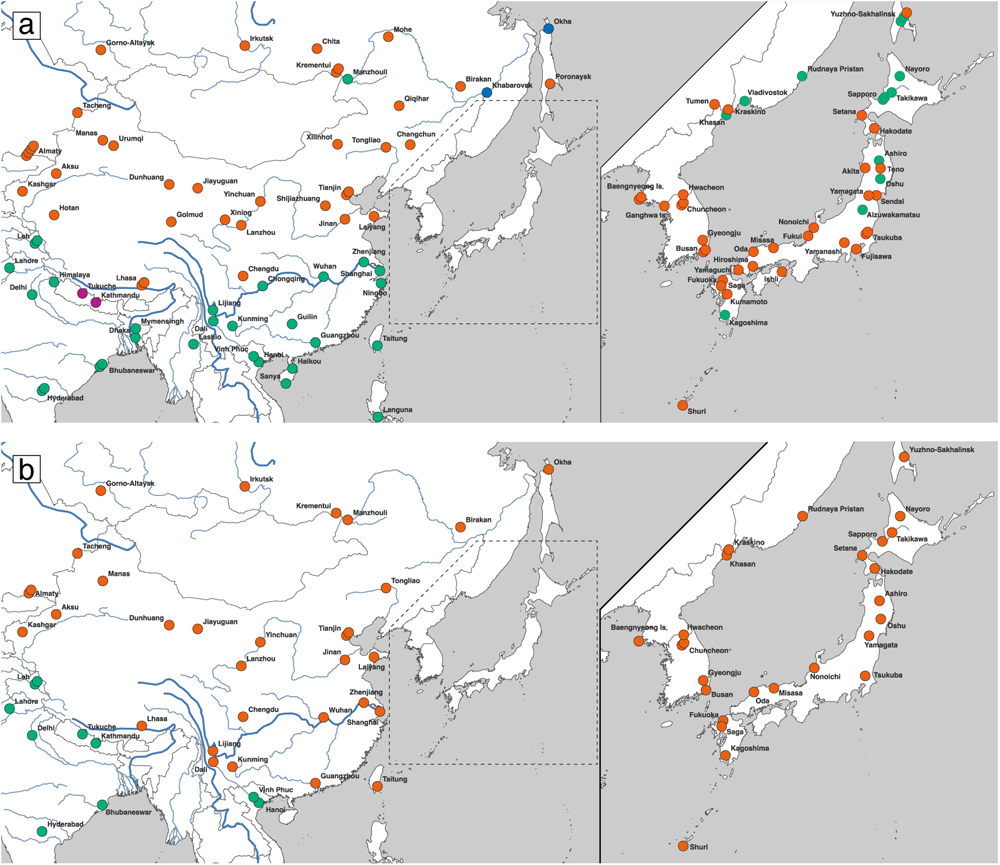
The geographic distribution of the subspecies *castaneus* and *musculus* haplotypes of mitochondrial and Y-chromosomal loci.

The upper panel (a) and lower panel (b) shows the mitochondrial and Y-chromosomal haplotypes, respectively. The colors of circles indicate subspecies haplotypes based on genotyping: *castaneus*-type (green), *domesticus*-type (blue), *musculus*-type (red), and geographically confined group of Nepalese-type (purple).

We hypothesized that the dominance of the *musculus*-type Y chromosome in East Asia is due to a distorted sex ratio caused by a dosage imbalance in the *Sly*/*Slx* genes; *Sly* and *Slx* are gametologs under X-Y sexual conflict and are highly duplicated genes (>100 copies). Previous studies have shown that *musculus*-type Y chromosomes encompass more *Sly* copies and the subspecies *musculus* have more copies of *Slx* on the X chromosome (Morgan and Pardo-Manuel de Villena 2017). Estimating the copy numbers of *Sly* and *Slx*, however, is challenging due to the presence of highly similar but distinct classes of repetitive sequences in the genome. To tackle this, we first aligned the paralogous sequences of the *Sly* and *Slx* loci present in the reference genome and used these alignments to reconstruct a phylogenetic tree. The reconstructed tree showed two clusters for *Slx*, which correspond to *Slx* (X1) and *Slxl1* (X2), and six clusters for *Sly* (Y1–Y6), with one cluster (Y1) being the most abundant in the reference genome and including the canonical *Sly* (Fig. 3a). We then estimated the copy numbers for each cluster based on the short-read depth of cluster-specific SNP alleles (see Methods). We observed that different subspecies-level Y haplotypes harbored different patterns of *Sly* clusters (Fig. 3b). The cluster Y5 was the major cluster in the *castaneus*-type haplotypes, while the cluster Y1 was the major cluster in the *domesticus*-type, and *musculus*-type Y haplotypes, respectively. Particularly, *castaneus*-and *domesticus*-type Y haplotypes had low *Sly* copy numbers, while *musculus*-type Y haplotypes had high copy numbers among the three haplogroups. In contrast, the copy number ratios of *Slx* and *Slxl1* were mostly consistent among samples. As shown in Fig. 3b and the previous study, *Sly* and *Slx* copy numbers were highly correlated (*P* < 10^−22^, Spearman’s rank correlation test) (Morgan and Pardo-Manuel de Villena 2017).

**Fig. 3:**
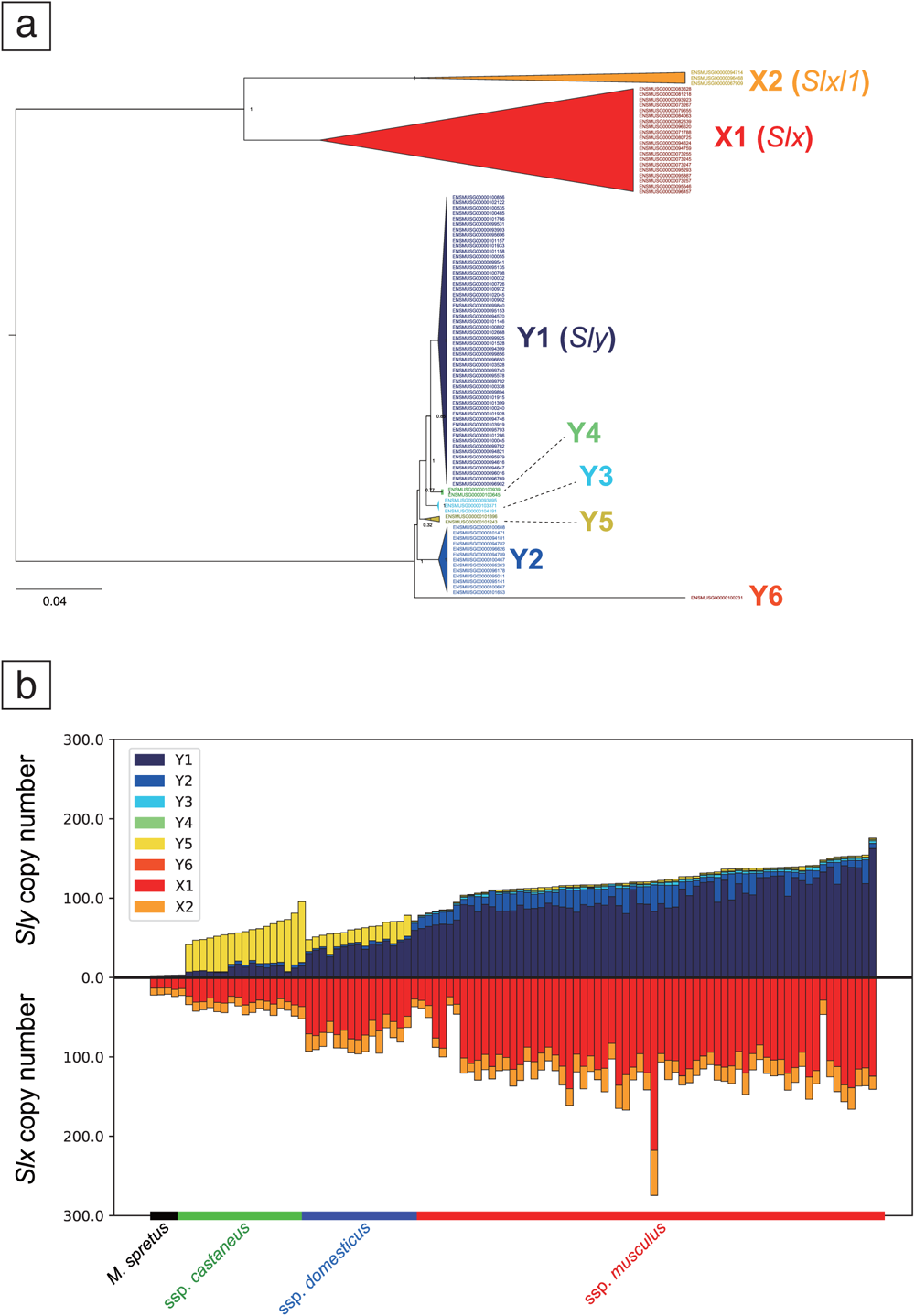
Copy number evolution of *Sly* and *Slx*.

The phylogenetic tree of *Sly*/*Slx* genes on the mouse reference genome, GRCm38 (a). Bootstrap values are described at each node. Estimated copy numbers of *Sly* and *Slx* for each sample (b). Different colors represent the copy numbers of different cluster. The samples were aligned according to the Y chromosome haplotypes first and sorted ascendingly by the total copy numbers of *Sly*.

In order to investigate selective forces shaping the sweep-like pattern of Y chromosome haplotypes in southern China, we performed Wright-Fisher computer simulations mimicking the introgression from the subspecies *musculus* population to subspecies *castaneus* populations in southern China. The model assumes that a small fraction of the subspecies *musculus* individuals with longer haplotypes of SD locus on the sex chromosomes (corresponding to *Sly* and *Slx*) continuously migrate to the subspecies *castaneus* population. An F1 hybrid between a *musculus-* type male and a *castaneus-*type female tends to have more male offspring than female offspring owing to the segregation distortion of X/Y chromosomes in a male germline. We also modeled loci responsible for partial hybrid incompatibility on the X chromosome (XHI loci). Detailed methods and results are documented in the Supplementary Note 1, Supplementary Figure 5, and Supplementary Table 2. Briefly, without assuming the mutations on copy numbers at the SD locus, the simulations showed that the introgressed Y chromosomes with longer SD haplotypes (corresponding to the *musculus-*type *Sly*/*Slx* haplotypes) quickly spread in the recipient population. The X chromosomes with longer SD haplotypes also eventually fix in the recipient population but the fixation of longer haplotypes was in general faster on the Y chromosomes than on the X chromosomes, because we assumed that SD occurs on male germline. The difference of fixation time between the Y and X chromosomes becomes longer when the effect of XHI is strong and the recombination between XHI loci and SD locus on the X chromosome is rare. When the effect of XHI is sufficiently strong, the introgression of X chromosomes with longer SD haplotypes are completely suppressed.

### Selection during the admixture in the Japanese population

To identify genomic regions affected by natural/sexual selection during intersubspecific admixture, we focused on samples from the Japanese archipelago and searched for genomic regions highly enriched in *castaneus*-ancestry haplotypes. Following the results presented in Fig. 1, we estimated the values of *α*, representing the proportion of *musculus*-like genomic components, for each non-overlapping 20-kb-long window of the autosomal genome.

To discern the neutral distribution of *castaneus*-enriched genomic blocks within the Japanese house mouse population, we employed the Fastsimcoal2 software. This aided in inferring demographic parameters from our sample set, which were used in estimating *α* (See Supplementary Note 2, Supplementary Figure 6, and Supplementary Table 3 for details). Using these parameters, we executed 100,000 coalescent simulations to ascertain a distribution of *α* under neutrality. From our results, we identified 4194 windows. These windows, which fell below the 5% threshold of the neutral distribution, encompassed the protein-coding regions of 842 distinct genes. A comprehensive list of these genes can be found in Supplementary Table 4.

We performed a functional enrichment analysis on the 842 candidate genes using MetaScape software (Zhou et al. 2019). A total of 20 gene categories were found to be enriched in the list (*p* values < 0.01), with the complete list of significantly overrepresented functional categories presented in Supplementary Figure 7. The enriched term list contained a variety of gene categories, but notably, immune-related gene categories such as immunoglobulin production (GO:002377), antibody-dependent cellular cytotoxicity (GO:0001788), and Herpes simplex virus 1 infection (mmu05168) were overrepresented. Indeed, our screening accurately identified the retroelement-like *Fv1* (Friend virus susceptibility protein 1) gene, an antiviral factor (Best et al. 1996) as an *castaneus*-ancestry enriched gene in Japan. Previous studies have demonstrated that the *Fv1^b^* allele, which lacks the 1.3-kb-length deletion commonly found in the subspecies *domesticus* and *musculus*, is observed with high frequency in the subspecies *castaneus* and the Japanese samples (Boso et al. 2021). We confirmed that the deletion started from chr4:147,870,288 (GRCm38) and had 1.2 kb in length, resulting in the truncation of C-terminal region of *Fv1*. The allele frequencies of deletion were 0.92, 0.64, and 0.16 in the subspecies *domesticus*, *musculus*, and *castaneus*, respectively. The allele frequency in the Japanese samples was 0.02, showing that almost no Japanese house mice harbor the deletion.

We also observed that 23 vomeronasal receptor genes (*Vmn1r* and *Vmn2r*), including one pseudogene, were present in the gene list. In the enrichment analysis, a category, detection of chemical stimulus involved in sensory perception (GO0050907) showed the strongest bias. These *castaneus*-enriched vomeronasal receptor genes were scattered across different olfactory/vomeronasal genomic clusters on the mouse genome, with most genes located in a large cluster on chromosome 7: region chr7:84,853,553-87,037,968, containing 17 olfactory and 15 vomeronasal receptors. Of the 23 vomeronasal receptor genes identified in our genomic scan, 12 were located in the cluster. To confirm that the estimated low *α* in the region was not due to errors in statistical inference, we computed *F*_ST_ between Japanese and subspecies *musculus* samples (*_F_*_ST-MUS/JPN_) and *F*_ST_ between Japanese samples and subspecies *castaneus* samples (*_F_*_ST-CAS/JPN_). The subspecies *castaneus* and *musculus* samples were assigned from the result of the ADMIXTURE analysis. We plotted window-averaged *F*_ST-CAS/JPN_ and *F*_ST-MUS/JPN_ across the region including chr7:84,853,553-87,037,968 (Fig. 4). As expected, outside of this region, *F*_ST-MUS/JPN_ values were consistently smaller than *F*_ST-CAS/JPN_, supporting the notion that Japanese samples were generally *musculus*-like. However, the pattern was completely reversed within the olfactory/vomeronasal clusters, indicating an unusual pattern of genetic admixture in this region. We also estimated genealogies around the target nonsynonymous SNVs of *Vmn2r* genes using RELATE software (Speidel et al. 2019). The genealogies encompassing the SNVs showed that the *castaneus*-type Japanese alleles coalesced with the other *castaneus*-type alleles after the split of the subspecies *castaneus* and *musculus*, which was estimated to occur around 200 kya in our data. The examples of nonsynonymous SNV segregation patterns and estimated genealogies are shown in Fig. 5.

**Fig. 4:**
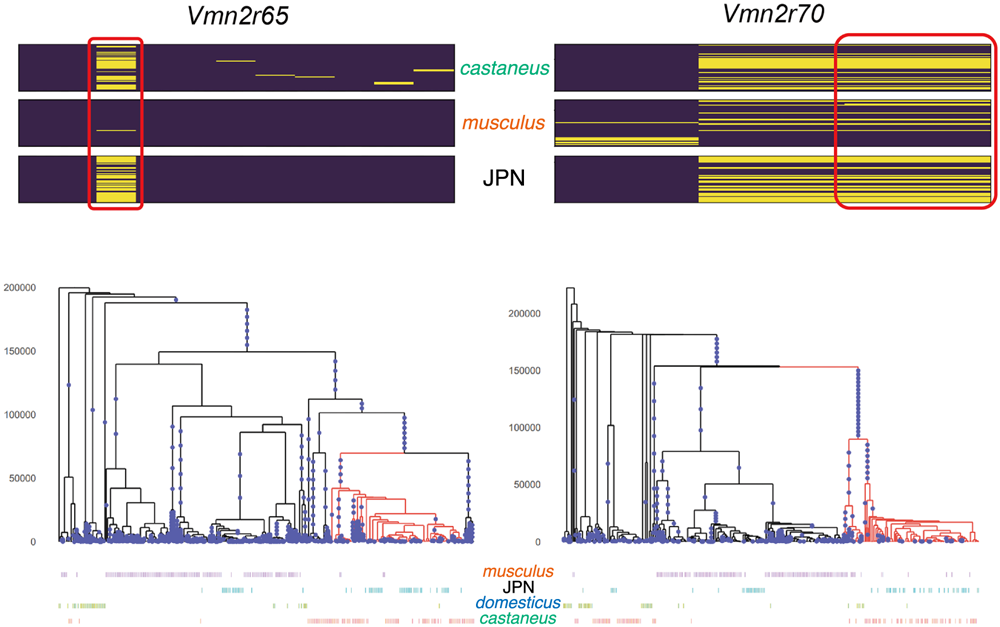
Haplotype structure of nonsynonymous SNVs and genealogies of wild house mouse.

**Fig. 5:**
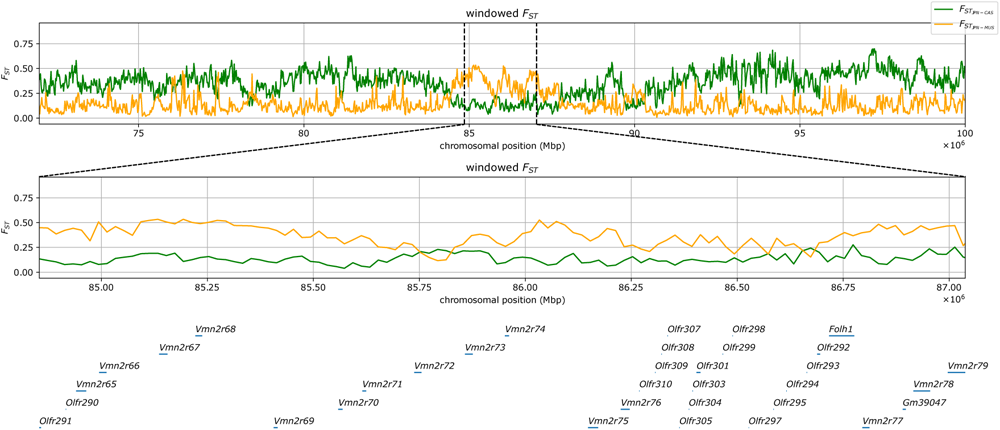
Distribution of *F*_ST_ in 20kb-length-windows on chromosome 7.

### Vmn2r65 and Vmn2r70

The upper panels show the segregation pattern of nonsynonymous SNVs. The sites in the red rectangles represent the focal sites that showed *castaneus*-ancestry enrichment in the Japanese (JPN) samples. The lower panels show the inferred genealogies around the focal sites. The branches with the derived mutations at the focal sites are colored in red. The labels of samples are shown at the bottom of the tree.

The yellow and green lines indicate *F*_ST_ between the Japanese and subspecies *musculus* samples and *F*_ST_ between the Japanese and subspecies *castaneus* samples, respectively. The lower panel shows the pattern in the olfactory/vomeronasal genes in this cluster (chr7:84,853,553-87,037,968, GRCm38).

## Discussion

### Genetic differentiation and inferred history of M. musculus in East Asia

The widespread coexistence of the subspecies *castaneus* and *musculus* in East Asia suggests that they formed a broad hybrid zone following their initial migration from their homeland. Previous studies have inferred migration scenarios primarily based on phylogeographical patterns of mitochondrial loci (Jing et al. 2014; Li et al. 2020). We present a combined view from autosomal, mitochondrial, X-chromosome, and Y-chromosome loci results in Supplementary Note 3.

### Spread of musculus-type Y chromosomes in East Asia

In this study we uncovered natural variation in Y chromosome of house mouse across Eurasia, identifying distinct clusters corresponding to major subspecies. Surprisingly, all Y chromosomes in East Asian samples were of the *musculus*-type, which could be explained by various mechanisms such as natural/sexual selection, male-biased migration, and genetic drift. Genomic conflict under the *Sly*/*Slx* system is likely a key factor, where increased *Sly* copy number promotes *musculus*-type Y chromosome spread in admixed populations like Japan and southern China. In Europe, hybridization reduces the selective advantage of *musculus*-type Y chromosomes due to male hybrid sterility, while *musculus*-type Y chromosomes have an advantage over *castaneus*-type Y chromosomes in interbreeding with subspecies *castaneus*. Our cluster-level analysis of *Sly* copy number variations revealed that *Sly* alleles, characterized by distinct amino acid sequences, expanded disparately across various Y chromosome lineages. These distinct sequences might confer unique functional attributes to each *Sly* allele. The question of whether these different amino acid sequences result in functional differentiation among *Sly* alleles remains a captivating question in future studies.

### Post-admixture natural selection in the Japanese archipelago

The Japanese samples showed high *castaneus*-ancestry in certain genomic regions despite their predominantly *musculus*-like genetic background. As demonstrated in previous studies (Lawal et al. 2021), genes involved in host defense mechanisms were significantly enriched in the outlier loci. In the case of humans, several studies have shown evidence of adaptive introgression from archaic to modern humans at immune-related loci (Racimo et al. 2015; Enard and Petrov 2018). For example, we identified *Irgm1* and *Irgm2*, which are GTPases involved in the interferon signaling pathway, in our candidate list. In particular, *Irgm2* has been shown to be responsible for defense against Toxoplasma, and numerous *castaneus*-like tightly-linked nonsynonymous SNVs were common in the Japanese samples (Supplementary Figure 8 and 9). In another example, the *castaneus*-ancestry enriched regions contained six cathepsin genes on chromosome 13. Two of them, *Ctsj* and *Ctsr*, are exclusive to adult placentas, and may be responsible for the process of viral infection, such as SARS-CoV-2 (Zhao et al. 2021). Although the roles of *Ctsj* and *Ctsr* in viral infection has to be demonstrated, maternal–fetal viral transmission might be a critical factor during the process of hybrid formation.

In addition to those genes, vomeronasal receptors exhibited a strongly biased pattern of segregation. These receptors are expressed in the vomeronasal organ and form a large receptor family involved in vomeronasal chemosensation (Wynn et al. 2012). They recognize a wide variety of chemical cues, such as pheromones from different sexes and predator odors (Dulac and Torello 2003). In some vomeronasal receptor genes, most Japanese samples harbor *castaneus*-type nonsynonymous alleles. Although certain vomeronasal receptor genes are highly differentiated between house mouse subspecies and could serve as markers for discrimination (Wynn et al. 2012), our genealogical analysis revealed that the target genes frequently crossed subspecies boundaries (Fig. 4). It is thus possible that some vomeronasal alleles become adaptive in other subspecies and quickly spread following introgression. Interestingly, frequent introgression of olfactory and other chemosensory receptor genes, but not particularly vomeronasal receptor genes, have been observed in the analysis between the subspecies *domesticus* and *musculus* (Staubach et al. 2012; Janousek et al. 2015).

The identified vomeronasal receptors were primarily clustered in a region on mouse chromosome 7, chr7:84,853,553-87,037,968. This entire ≈2Mb region displayed a strongly biased pattern of *castaneus*-ancestry (Fig. 5). Interestingly, previous extensive studies revealed that *Vmn2r* genes in this cluster respond to cues from conspecific mating partners (Isogai et al. 2011). Detailed studies, such as gene replacement experiments, will elucidate how different vomeronasal receptor alleles contribute to the formation of natural house mouse hybrids in the future.

Altogether, we have elucidated the genomic landscape of wild house mice in East Asia, demonstrating a deep connection between human and mouse genetic histories. In East Asian house mouse populations, which experienced secondary contact after the subspecies split, we have inferred the effects of natural/sexual selection and genomic conflict from the observed pattern of genetic structure at various genomic loci. These findings will pave the way for future studies on genetic admixture between different house mouse subspecies and provide insights into the genetic architecture of hybrid genomes.

## Methods

### Genomic samples

We have sequenced the whole genomes of 37 *M. musculus* samples, with mitochondrial DNA sequences published in previous studies (Moriwaki et al. 1986; Suzuki et al. 2004; Terashima et al. 2006; Nunome et al. 2010; Suzuki et al. 2013; Kodama et al. 2015; Kuwayama et al. 2017). We combined this data with the global sample dataset used in our prior research (Fujiwara et al. 2022a). A detailed list of these samples can be found in Supplementary Table 1. In total, our analysis included 163 *M. musculus* and 7 *M. spretus* samples.

### Mapping and genotype calling

We sequenced the genomes of the 37 samples using the DNBSEQ platform (100-or 150-bp-length paired-end). These newly sequenced whole genome samples were the samples used primarily for mitochondrial analysis in previous studies. The methodology for quality checks, mapping reads, and genotype calling follows the same approach as in our previous study (Fujiwara et al. 2022a). We used the GRCm38 (mm10) assembly as the autosomal reference, and the RefSeq sequences NC_005089 and NC_205952 as the mitochondrial references for *M. musculus* and *M. spretus*, respectively. To determine the sexes of the 37 samples, we counted the number of mapped reads to sex chromosomes as described in our previous studies (Fujiwara et al. 2022a). The determined sexes for these samples can be found in Supplementary Table 1.

### Autosome re-sequencing analysis

We conducted a principal component analysis on 163 *M. musculus* samples to investigate population structure. This analysis was performed using smartpca from the Eigensoft software with default parameters, except that outliers were not excluded (Patterson et al. 2006).

To further analyze population structure and admixture, we employed ADMIXTURE with the number of clusters *K* = 3 for the 163 *M. musculus* samples. Before admixture inference, we pruned SNPs under linkage disequilibrium (LD) using PLINK with the option “--indep-pairwise 50 5 0.5”. In downstream analyses, the majority of ancestral proportions determined by ADMIXTURE analysis define the subspecies of each sample.

To quantify the fraction of genomes contributed by the subspecies *musculus* (α) in the East Asian samples, we estimated *f*4-ratio statistics (Patterson et al. 2012). The estimation of *f*_4_ statistics was carried out using the “patterson_d” function in the Scikit-allel Python package. Assuming the source of *musculus* genomic components was more closely related to the Korean *musculus* samples than to Kazakh *musculus* samples, we estimated the α value for sample X as equation (1):

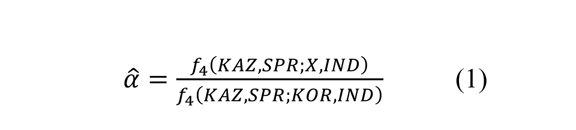

, where KAZ, SPR, KOR, and IND represent the samples from Kazakhstan (5 samples), *M*. *spretus* (7 samples), Korea (9 samples), and India (2 samples from Leh), respectively. The model is also presented in Supplementary Figure 6. In principle, *α* is equal to or smaller than 1, but it may slightly exceed 1 because of statistical fluctuation. When *α* >1, we therefore set the *α* to 1. Additionally, we estimated the values of α for each 20kb-length window for all Japanese samples. To remove unreliably estimated values, we excluded windows with fewer than 100 SNVs. Both the denominator and numerator of the right side of equation 1 are expected to be positive, considering that the Japanese and Korean samples are genetically closer to the subspecies *musculus* (KAZ samples) than to the subspecies *castaneus* (IND) samples. However, a very small fraction of windows showed negative denominators and/or numerators due to statistical errors, introgression, or ancestral polymorphisms. Such windows were filtered out from the following analysis. *F*_ST_ was estimated using the “hudson_fst” function in Scikit-allel.

### Mitochondrial genome assembly

Complete mitochondrial sequences of all 37 samples were *de novo* assembled using GetOrganelle software (Jin et al. 2020) with maximum extension rounds set to 10 and *k*-mer parameters set to “21, 45, 65, 85, 105, 127”. The qualities of assembled mitochondrial genomes were visually checked using Bandage software (Wick et al. 2015). All sample sequences were aligned using Clustal Omega with the “--auto” option. All D-Loop regions and gapped sites were removed from the alignments. A maximum-likelihood phylogenetic tree was constructed using IQ-TREE2 software (Nguyen et al. 2015) with the bootstrapping of 1000 replications. Before constructing a phylogenetic tree, we performed ModelFinder software (Kalyaanamoorthy et al. 2017) implemented in IQ-TREE2 to determine the best substitution models. According to ModelFinder, “TPM2+F+I+G4” was chosen as the best substitution model using the Bayesian Information Criterion.

### Y chromosome re-sequencing

We re-sequenced the Y chromosomes using raw read data obtained by whole genome sequencing. The list of male samples that were used for the Y genotyping is shown in Supplementary Table 1. The SNVs and Indels were called by GATK4 (McKenna et al. 2010) HaplotypeCaller with “-ERC GVCF” option to obtain genomic variant calling files (gVCF). All sample gVCF were then genotyped by GATK4 GenotypeGVCFs with “-all-sites” option. We used GenMap software to calculate mappability scores with “-K 30” and “-E 2” options, and filtered out the positions with mappability score <1.

For the phylogenetic analysis, we restricted our analysis to the short arm of this Y chromosome (the first 3.4 Mb length), because the vast majority of the long arm of Y is composed of amplicon sequences. We also excluded SNVs that were labeled as “LowQual” by the GATK4 HaplotypeCaller and those with read depths ≤3. The obtained VCF file of the Y chromosome was converted to Fasta, and Nexus files. The maximum likelihood phylogenetic tree of the Y chromosome was constructed by IQ-TREE software (Nguyen et al. 2015) with the bootstrapping of 1000 replications. According to ModelFinder, “TIM2+F+R2” was chosen as the best substitution model using Bayesian Information Criterion.

### Copy number estimation of Sly/Slx

We downloaded all the paralogous sequences of *Sly* (ENSMUSG00000101155) and *Slx* (ENSMUSG00000095063) on the sex chromosomes, including introns, from Ensembl Release 102. The sequences were then aligned using MAFFT (Katoh et al. 2002; Katoh and Standley 2013). Following the alignment, we reconstructed a phylogenetic tree using MEGA X software (Kumar et al. 2018), employing a maximum likelihood method with the GTR (General Time Reversible) model. From this alignment, we identified tag-SNP sites with cluster-specific alleles.

We started by mapping the short reads from samples to either *Sly* or *Slx* genomic sequences, incorporating 1000-bp-length flanking sequences for both ends. The copy numbers for each cluster were then estimated by counting the depth of cluster-specific alleles at the tag-SNP sites, using the pileup format files generated from samtools mpileup command. To filter out results from repetitive sequences that might cloud our analysis, we focused solely on sites with a mappability score of 1, calculated using autosomal sequences and either *Sly* or *Slx* sequences. To ensure the accuracy and relevance of our results, a filter was applied to exclude sites that exhibited a depth smaller than 1/3 or larger than three times the median. Finally, we estimated the copy number for each sample from the mean depth, which was then divided by half the average depth of the whole genome for the sample.

### Deletion detection from sequence data

DELLY (v.1.1.6) was used for genotyping retroelement-like *Fv1* (Friend virus susceptibility protein 1) gene deletion. Since location of *Fv1* gene positioned between chr4:147,868,979– 147,870,358 in chromosome 4, we restrict regions for detecting deletions between chr4:145,000,000–150,000,000 to speed up calculations.

### Construction of genome-wide genealogies using RELATE

We estimated the genome-wide genealogies using RELATE v.1.1.9 (Speidel et al. 2019). The input files were converted from variant call format (.vcf) to haplotype format (.haps) using the RelateFileFormats command with the “--mode ConvertFromVcf” option implemented in the RELATE package. Non-biallelic SNVs were removed from the analysis, and the *M. spretus* genome was used to assign derived mutations in *M. musculus*. A germline mutation rate of 5.7 × 10^−9^ per bp per generation (Milholland et al. 2017) and a generation time of 1 year, and an initial effective population size parameter of 120,000 were used for estimation. We then used the EstimatePopulationSize.sh script in the RELATE package to re-estimate the effective population size changes over time. We set the threshold to remove uninformative trees at 0.5.

## Data access

Supplementary material is available online. All raw short-read sequencing data generated in this study have been submitted to the DNA Data Bank of Japan (DDBJ) DDBJ Sequence Read Archive (DRA) database under accession number PRJDB16017. All complete mitochondrial sequence generated in this study have been also submitted to the DDBJ with accession number LC772928-LC772964. The data sets, parameterization files, and scripts required to reproduce the analysis are deposited in the Dryad digital repository (https://datadryad.org/) under doi: [https://datadryad.org/stash/share/AxCAUN6Te8zqxpLB9C87BbtTmjJbBaBHBPdNcGZsqss].

## Acknowledgments

We would like to express our sincere gratitude to the late Dr. Kazuo Moriwaki for collecting valuable wild house mouse samples from around the world. We thank the members of the Yaponesian Genome project funded by the MEXT KAKENHI, for their valuable comments and feedback to our research. This work was supported by MEXT KAKENHI (grant 18H05511 and 23H04846 to N. O. and grant 18H05508 to H. S.)

