## Supplementary for "Inference of Selective Force on House Mouse Genomes during Secondary Contact in East Asia"

Table of Contents:


1 **Supplementary Note 1**: Computer simulations for the introgression of X-chromosomal, Y-chromosomal, and mitochondrial genomes under sex-ratio distortion

4 **Supplementary Note 2**: Demographic inference and computer simulations for admixture

6 **Supplementary Note 3**: Dispersal model of *M. musculus* in East Asia

10 **Supplementary Figure 1**: Principal component analysis (PCA) of wild house mouse using 163 *Mus musculus* samples

11 **Supplementary Figure 2**: ADMIXTURE plot of *Mus musculus* samples using autosome data

12 **Supplementary Figure 3**: The maximum likelihood tree of mitochondrial genomes

13 **Supplementary Figure 4**: The maximum likelihood tree of the short arm of the Y chromosomes

14 **Supplementary Figure 5**: Trajectories of allele frequency of introgressed alleles

15 **Supplementary Figure 6**: The demographic model we used to estimate the population genetic parameters

16 **Supplementary Figure 7**: Functional enrichment analysis of genes with *castaneus*-ancestry bias

17 **Supplementary Figure 8**: Haplotype structure of nonsynonymous SNVs and genealogies of wild house mouse *Irgm1*

18 **Supplementary Figure 9**: Haplotype structure of nonsynonymous SNVs and genealogies of wild house mouse *Irgm2*

19 **Supplementary Table 1**: List of samples used in this study

21 **Supplementary Table 2**: Allele frequency of introgressed alleles with different *r*

22 **Supplementary Table 3**: Estimated population genetic parameters and confidence intervals

23 **Supplementary Table 4**: List of genes in the *castaneus*-enriched 20-kb-length windows

### Supplementery Note 1

*Computer simulations for the introgression of X-chromosomal, Y-chromosomal, and mitochondrial genomes under sex-ratio distortion*

#### Model

We modeled a Wright-Fisher diploid population A of size $N$, which consists of males and females with the XY sex-determination system. Population A receives migrants from another population B at a rate of $m$ (fraction of migrants in the recipient population) per generation. Assuming that populations A and B have already well differentiated genetically, for each individual in population A, we consider five key genetic parameters: sex, a genomic fraction derived from population B ($G_A$), genotype at sex-ratio distortion (SD) locus on X ($G_{XSD}$) and/or Y ($G_{YSD}$) chromosomes, genotype at X-chromosomal hybrid incompatibility (XHI) locus ($G_{XHI}$), and mitochondrial genotype ($G_M$). We assumed that SD and XHI loci are recombined with a rate of $r$ per generation. In the following definitions, we assigned the value 0 and 1 to the original alleles in populations A and B, respectively.

We assumed that the autosomal genome would experience a sufficient amount of recombination. The feature of the offspring's genome, $G_A$, is scored by an average of parental values. $G_A$ of the F1 hybrid between an individual from unadmixed populations A and B would thus become 0.5, and $G_A$ of the backcrossed hybrid in population A would be 0.25. The genetic distance between parental genomes at autosomal loci, $d_{AA}$, was defined as the absolute difference between $G_A$ values of a father and a mother. Similarly, the genetic distance between autosomes and X chromosomes, $d_{AX}$, was defined as the absolute difference between $G_A$ and $G_{XHI}$. Since a female offspring has two X chromosomes, $G_{XHI}$ in females was averaged over the two chromosomes. The genetic distance between autosomal and mitochondrial genomes, $d_{AM}$, was defined in a similar way.

The SD locus is located on both X and Y chromosomes. We assumed that the SD locus has copy number polymorphisms and that allele type 1 in population B has a higher copy number than allele type 0 in population A. In a male germline, a sex chromosome that has an SD allele with a higher copy number than the other sex chromosome is assumed to have a twice higher probability of transmission to the offspring. When the allele on the Y chromosome is 1 and the allele on the X chromosome is 0, the male-to-female ratio of offspring would be 2.

The effect of hybrid incompatibility was evaluated for each reproduced offspring. Assuming the multiplicative effect of hybrid incompatibility between autosomal genomes, between the X chromosome and autosomal genomes, and between the mitochondrial and autosomal genomes, the fitness of offspring, $w$, is defined as equation (1):

$$w = exp[-\alpha d_{AA} - \beta d_{AX} - \gamma d_{AM}] \quad (1)$$

, where $\alpha$, $\beta$, and $\gamma$ represent amplitude factors for each effect.

### Simulations

We developed a Python script for an individual-based population genetics simulation, which consists of $N_m$ males and $N_f$ females in population A. $N_m$ and $N_f$ may not always be equal, but their sum, $N$, remains constant. Each generation in the simulation has two phases.

In the first phase, recombination between each XHI locus and SD locus on the X chromosomes in females is randomly assigned. Random variables are drawn from the Poisson distribution with a mean of $2rN_f$, and the specified number of recombination events is randomly allocated to the pool of X chromosomes in females. If mutations are considered, they are assigned to each genome during this phase in a similar manner.

In the second phase, mating pairs are randomly selected from population A. However, there is a probability of $m$ that one mate will be replaced by a migrant from population B. The offspring's sex ratio is determined by the father's genotype, as described in the previous section. After selecting mating pairs, the offspring's fitness is evaluated. A random variable is drawn from a uniform distribution U(0, 1). If the random variable is higher than $w$, the individual is discarded, and a new mating pair is chosen until $N$ offspring are produced for the next generation. This process is repeated for an adequate number of generations.

### Results

We performed simulations using various parameter settings and present results with $N$ = 1,000 for up to 100 (0.1$N$) generations. The number of generations is sufficient, considering that house mice have a population size on the order of 100,000–1,000,000, and we take into account the admixture events occurring around these 10,000 (0.1$N$) generations. We also fixed the migration rate $m$ at 0.0001. We initially ignored the recombination between the SD and XHI loci ($r$ = 0) and discuss its effect in a later part.

We first consider the case without any hybrid incompatibility ($\alpha$ = 0, $\beta$ = 0, and $\gamma$ = 0). The fixation rates of introgressed X, Y, and mitochondrial genomes after 0.1$N$ generations were evaluated using the results of 1,000 instances of simulations. Within 0.1$N$ generations, the fixation of introgressed X and Y chromosomes consistently occurred (1,000 instances). On the other hand, introgressed mitochondria did not fix in the recipient population, because the length of 0.1$N$ generations was not sufficiently long for neutral loci to be fixed even with a smaller effective population size at the mitochondrial locus. The result is shown in Supplementary Table 2. Trajectories of allele frequencies in population A are shown in Supplementary Fig. 5a. Interestingly, the fixation of introgressed X chromosomes usually occurred after the fixation of introgressed Y chromosomes. This would be because the spread of the introgressed alleles is mainly enhanced by the segregation distortion in male gametes. We next incorporated the effect of hybrid incompatibility due to the mismatch in the autosomal genomes ($\alpha = 1, \beta = 0$, and $\gamma = 0$). The parameter set corresponds to the fitness reduction of F1 hybrids to 37%. We found that the level of hybrid incompatibility would certainly reduce the rate of introgression, but the fixation probability of introgressed alleles was still high with the selective advantage of introgressed X and Y chromosomes (Supplementary Table 2). As expected, the rate of fixation was reduced when $\alpha$ was much greater than 1 (Supplementary Table 2). Trajectories of allele frequencies in population A are shown in Supplementary Fig. 5b. The effect of the XHI locus was incorporated in the next case ($\alpha = 1$, $\beta = 1$, and $\gamma = 0$). The X chromosomal effect for hybrid incompatibility greatly reduced the fixation rate of introgressed X chromosomes but did not change the fixation rate of introgressed Y chromosomes (Supplementary Table 2). Trajectories of allele frequencies in population A are shown in Supplementary Fig. 5c. We also incorporated the effect of mitonuclear incompatibility, where introgressed mitochondrial genomes are incompatible with the genomic background of recipients. Trajectories under the case $\alpha = 1$, $\beta = 1$, and $\gamma = 1$ are shown in Supplementary Fig. 5d, and fixation rates of introgressed alleles are presented in Supplementary Table 2.

The recombination rate between the SD and XHI loci, denoted as $r$, is a crucial factor in determining the fixation rate of introgressed SD alleles on the X chromosome. The simulation results with different levels of $r$, specifically $r = 0.1$, are presented in Supplementary Fig. 5. A moderate level of recombination readily broke up the linkage between SD and XHI loci, allowing introgressed SD alleles to spread rapidly within the recipient population. The effect of recombination was mitigated with a stronger X-chromosomal effect in hybrid incompatibility, as well as a stronger mitonuclear incompatibility.

Supplementery Note 2

*Demographic inference and computer simulations for admixture*

Demographic parameter estimation

In order to illustrate the distribution of *castaneus*-enriched genomic blocks in the Japanese population under neutrality, we inferred demographic parameters for the five population samples (*M. spretus*: SPR, Kazakhstan: KAZ, Korea: KOR, India Leh: IND, and Japan: JPN) using Fastsimcoal2 software (Excoffier et al. 2013). The total number of samples used for the analysis was 52. The model is presented in Supplementary Fig. 6. For simplicity, we assumed that each population size remained constant unless a population split or admixture occured, that migration rates were symmetric, and that a single pulse admixture event took place during the establishment of the Japanese population. We also assumed that the admixture forming the Japanese population occurred 3,000 generation ago, with the effects of this assumption discussed later in the text.

We generated joint site frequency spectrum data with minor allele frequencies for the five populations using the same samples employed in the calculation of the $f_4$ ratio ($\alpha$), which estimates the genomic proportion of *musculus*-derived alleles in the Japanese samples. In addition to the mappability filter, we also filtered our SNVs using the threshold of “ExcessHet>30”, marked by GATK.

Due to the limitations of our genotyping pipeline, it didn't specify the genomic sites that were monomorphic non-variant sites. To address this, we estimated the number of these sites. This estimation was essential to scale the parameters accurately. Our mappability mask identified 1,764,828,698 autosomal sites that passed the filtering process. Out of the 52 samples, we opted for the sites where all samples had been genotyped, leading to a reduction in polymorphic sites by a factor of 0.878. In addition, based on the H1KG project, we postulated that 93% of the autosomal genomes were accessible (http://ftp.1000genomes.ebi.ac.uk/). By combining this data, the total analyzed sites amounted to 1,440,533,483.

We assumed mutation rate of $5.9\times10^{-9}$ and generation time of 1 year, and repeated the maximum-likelihood estimation procedure 300 times, selecting the parameter sets that demonstrated the highest likelihood. We also conducted parametric bootstrap resampling 100 times to capture a 95% confidence interval. Details on the estimated population parameters can be found in Supplementary Table 3.

Computer simulations

We generated 20-kbp-length genomic fragments corresponding to the samples using the maximum-likelihood estimators in Supplementary Table 3, with a recombination rate of $1\times10^{-8}$ per site per generation, and estimated $\alpha$ 100,000 times. We assigned *p*-values to observed data using this distribution. When we did not fix the admixture timing for the Japanese population and estimated the timing using Fastsimcoal2, we obtained a slightly earlier admixture event estimate of >5,000 generation ago, which was much older than the assumed value. We are uncertain whether this deviation is due to our simplified model and assumptions; however, the older admixture event causes the distribution of α to skew toward higher values, making our test more conservative.

### Supplementary Note 3

*Dispersal model of M. musculus in East Asia*

The subspecies *musculus* reached East Asia through the northern side of the Himalayas, similar to Paleolithic human migrations from central/western Eurasia to East Asia (Osada and Kawai 2021). Its migration coincided with the transmission of wheats in the Late Neolithic period and foxtail and broomcorn millets in the Early to Middle Neolithic periods (Betts et al. 2014; Leipe et al. 2019). The migration route of the subspecies *castaneus* from North India to southern China is less clear, but it is likely that this migration was associated with rice cultivation. Whether japonica rice (*Oryza sativa japonica*) and indica rice (*O. s. indica*) were established independently remains controversial, but they experienced considerable gene flow after the initial domestication event, suggesting that there was human activity linking the two regions during the Early Neolithic period (Huang et al. 2012; Choi et al. 2017). The spread of the subspecies *castaneus* in Chinese and Southeast Asian regions was probably caused by the spread of rice cultivation.

Archaeological findings highlight three primary Neolithic cultural hubs in East Asia: the Yellow River basin and West Liao River basin, centered around with foxtail and broomcorn millet farming, and the Yangtze River basin, known for its rice cultivation. Recent ancient human genome sequencing has revealed that genetic differentiation between the people of these centers was much stronger than that of modern Chinese human populations (Ning et al. 2020; Yang et al. 2020). The current distribution of the subspecies *musculus* and *castaneus* corresponds to the genetic makeup of ancient humans and is associated with the spread of certain types of crop cultivation in East Asia. However, the degree of admixture between the subspecies *musculus* and *castaneus* appears to be much smaller than that between human populations in northern and southern China at non-Y chromosome loci, reflecting the limited mobility of house mice.

*Differentiation of M. musculus in the Japanese archipelago*

Both our previous and current analyses showed that the genomic components of the subspecies *musculus* are predominant in samples from the Japanese archipelago (Fujiwara et al. 2022). The strong genetic relatedness of Japanese and Korean samples at all genomic loci strongly supports the idea that the subspecies *musculus* was introduced to the Japanese archipelago by rice farmers with irrigation facilities (Yayoi people) who migrated through the Korean Peninsula beginning in 1,000 BCE (Fujio 2017). However, a recent study analyzing seed impressions in pottery revealed that early migrants from the Korean Peninsula to the Japanese archipelago during the Final Jomon and Initial Yayoi periods used both rice and millet (Endo and Leipe 2022). This suggests that the subspecies *musculus* may have reached the Japanese archipelago alongside millet farming, rather than rice farming. This is especially plausible given that the expansion of the subspecies *musculus* in Northern China is linked with millet farming.

Given that 10–20% of the modern Japanese genome is derived from continental migrants, human migration from continental Asia to the Japanese archipelago was intense and continuous (Hanihara 1991; Kanzawa-Kiriyama et al. 2019; Cooke et al. 2021; Osada and Kawai 2021). Considering that all *musculus*-type mitochondrial and Y-chromosome haplotypes in the Japanese samples are derived from one or two haplotypes from the Korean *musculus* lineage, it is likely that there were a limited number of subspecies *musculus* migrants. The subsequent population explosion of the subspecies *musculus* indicates its success in the rice-crop-oriented society in Japan (Yonekawa et al. 1988).

Our analysis reveals a clear pattern of differentiation along the coastline of the Japanese archipelago. Recent large-scale genome analyses of modern Japanese have not shown such a distinct trend (Watanabe et al. 2021). We propose that this pattern of differentiation was shaped by shipping trades. In the 18th century, Japan developed highly organized shipping trade routes along the Sea of Japan and Pacific Ocean coastlines. Ship travel may have facilitated the spread of the genetic component of *castaneus* from northern Japan to the Sea of Japan coast.

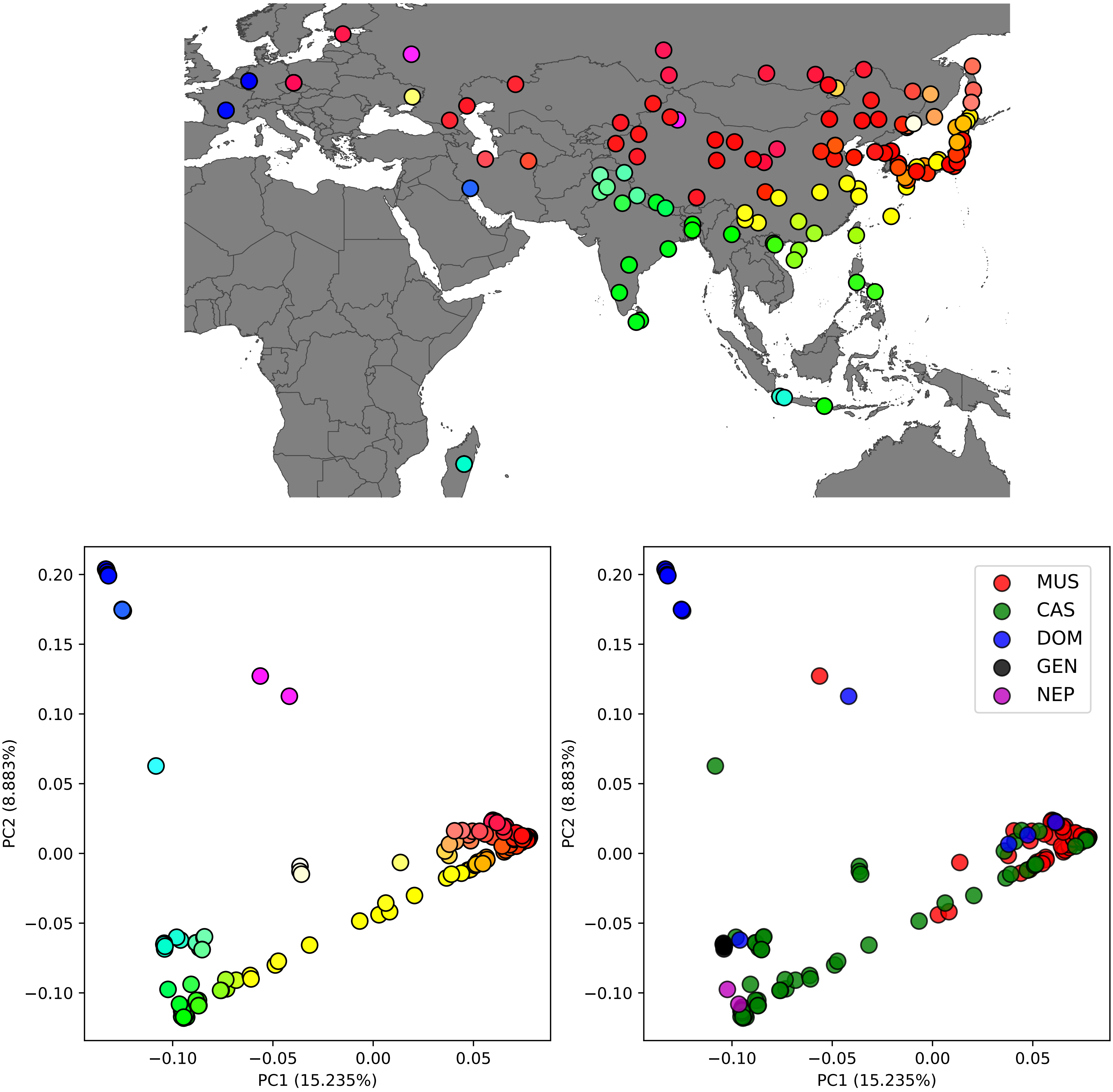


**Supplementary Figure 1.**

Principal component analysis (PCA) of wild house mouse using 163 *Mus musculus* samples. **(a)** Geographic map of the samples collected used in this study. Each circle represents an individual, and the color of the circle corresponds to the color used in panel (b). **(b)** Plot results of principal component analysis using autosomes of wild house mice. Red, green, and blue circles correspond to *musculus*, *castaneus*, and *domesticus* genetic components, respectively. The circles showing the intermediate color of each genetic component indicate the hybridization of each genetic component. The proportion of variance for each principal component (PC) is shown in each axis labels. **(c)** PCA results shown in panel (b) with mitochondrial haplogroup labels. The labels with MUS, CAS, and DOM represent the *musculus*, *castaneus*, and *domesticus* haplogroups, respectively. In addition, GEN and NEP represents distinct haplogroup in Madagascar and distinct haplogroup in Nepal, respectively.

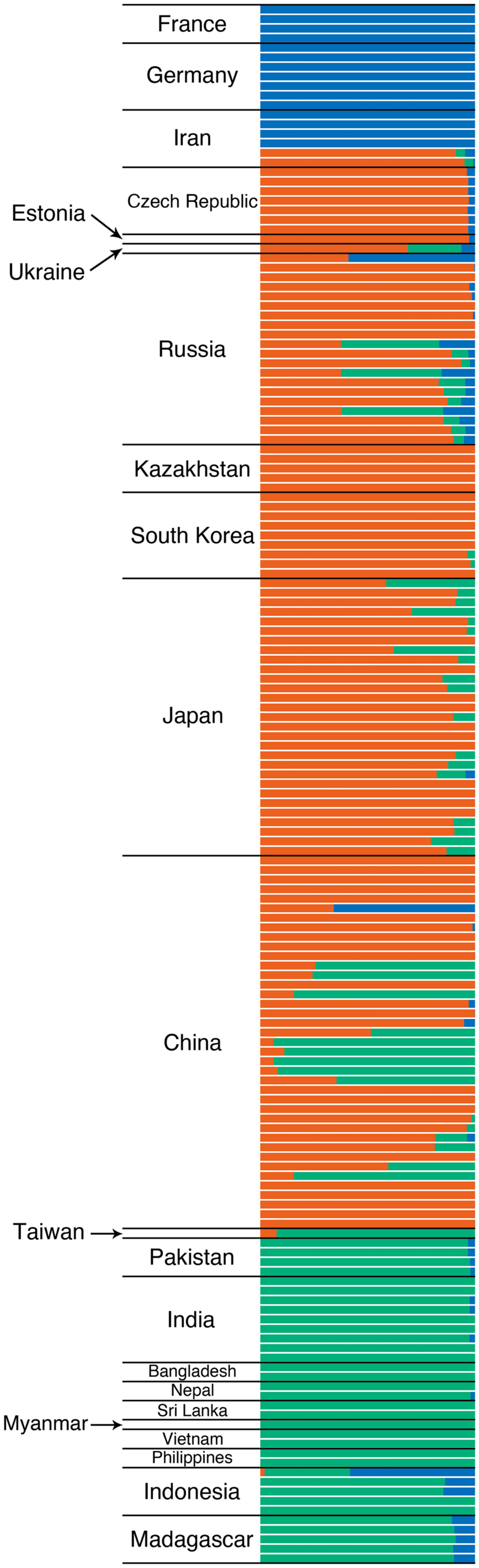


**Supplementary Figure 2**.

ADMIXTURE plot of *Mus musculus* samples using autosome data. Each bar represents each individual, and the color ratio represents the proportion of genetic elements. The results are shown for $K$=3, where red, green and blue colors represent the genetic elements of *musculus*, *castaneus* and *domesticus*, respectively.

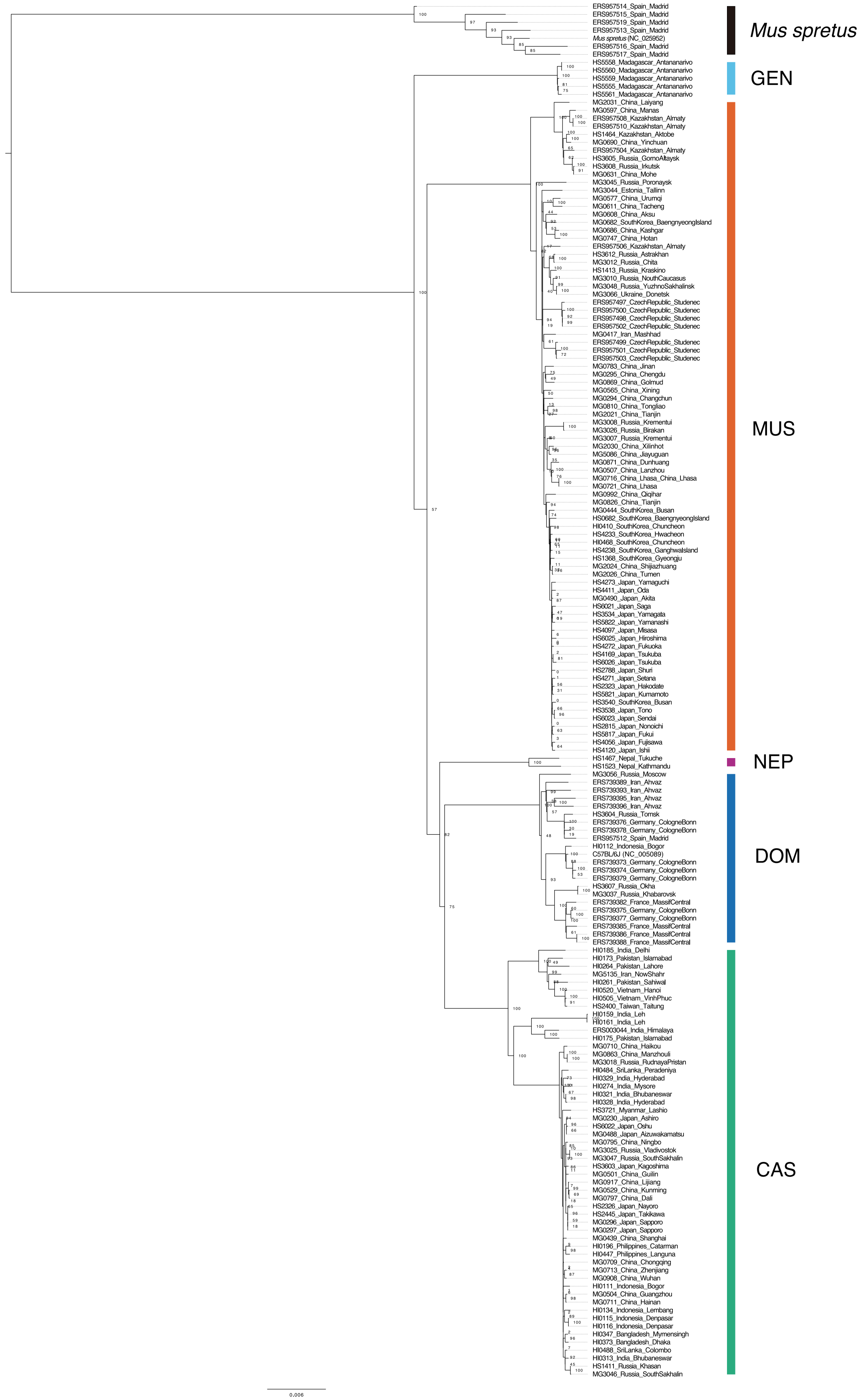


**Supplementary Figure 3.**

The maximum likelihood tree of mitochondrial genomes. The phylogenetic tree contains the mitochondrial sequences of 170 samples used in this study and additionally includes the mitochondrial reference sequences of *Mus spretus* (NC_025952) and *Mus musculus* (C57BL/6J strain, NC_005089). The substitution model (TPM2+F+I+G4) was determined as the best fit model by ModelFinder implemented in IQ-TREE2. Each captioned colored line indicates the corresponding clade of *Mus musculus* subspecies-derived haplotypes. The node labels exhibit support assigned by bootstrapping for 1000 replications. The labels with CAS, DOM, MUS represent *castaneus, domesticus, and musculus* haplogroups, respectively. GEN and NEP labels show distinct haplogroups in Madagascar and Nepal, respectively.

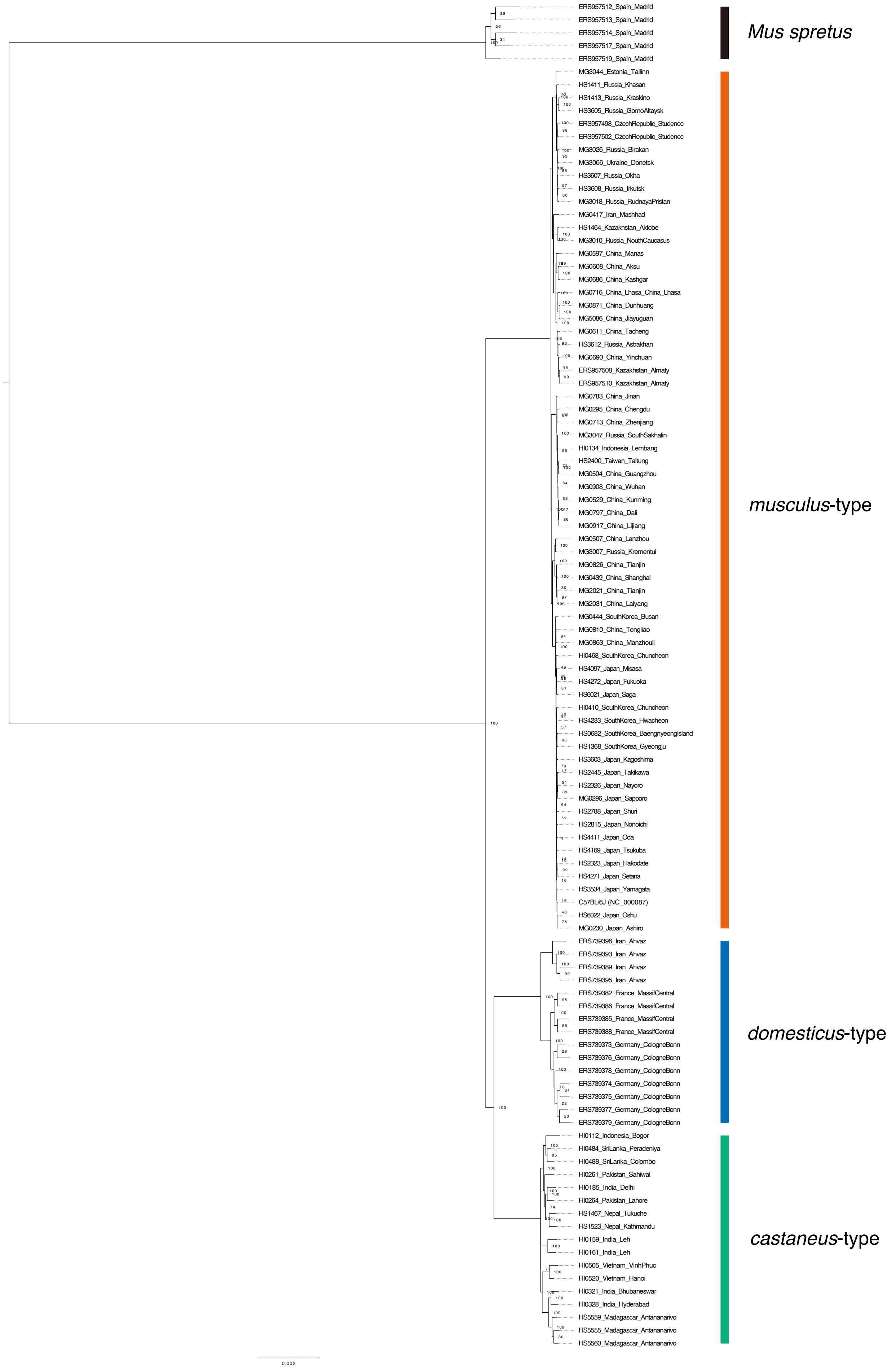


**Supplementary Figure 4.**

The maximum likelihood tree of the short arm of the Y chromosome. The phylogenetic tree contains the Y chromosome sequences of 103 male samples used in this study and additionally includes the Y chromosome reference sequence of *Mus musculus* (C57BL/6J strain, NC_000087). The substitution model (TIM2+F+R2) was determined as the best fit model by ModelFinder implemented in IQTREE2. Each captioned colored line indicates the corresponding clade of *Mus musculus* subspecies-derived haplotype. The node labels exhibit support assigned by bootstrapping for 1000 replications.

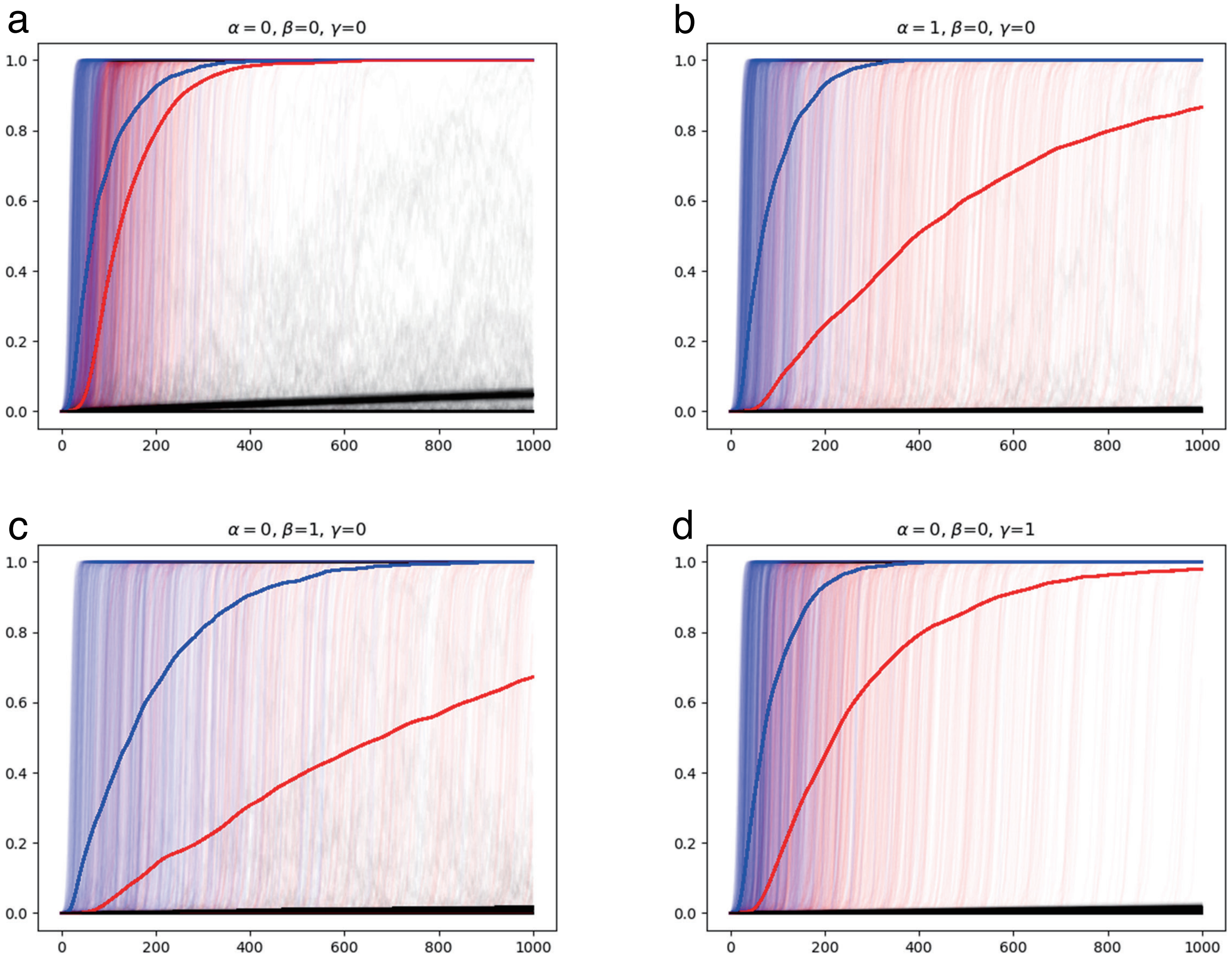


**Supplementary Figure 5.**
Trajectories of allele frequency of introgressed alleles. The red, blue, and black lines show the allele frequency at X-chromosomal, Y-chromosomal, and mitochondrial genomes, respectively. The thin lines represent the trajectories of single simulation and the thick lines show the average allele frequencies among 1000 trials. The parameter setting of each panel is shown in the top of plot. See Supplementary Note 1 for detailed methods and results.

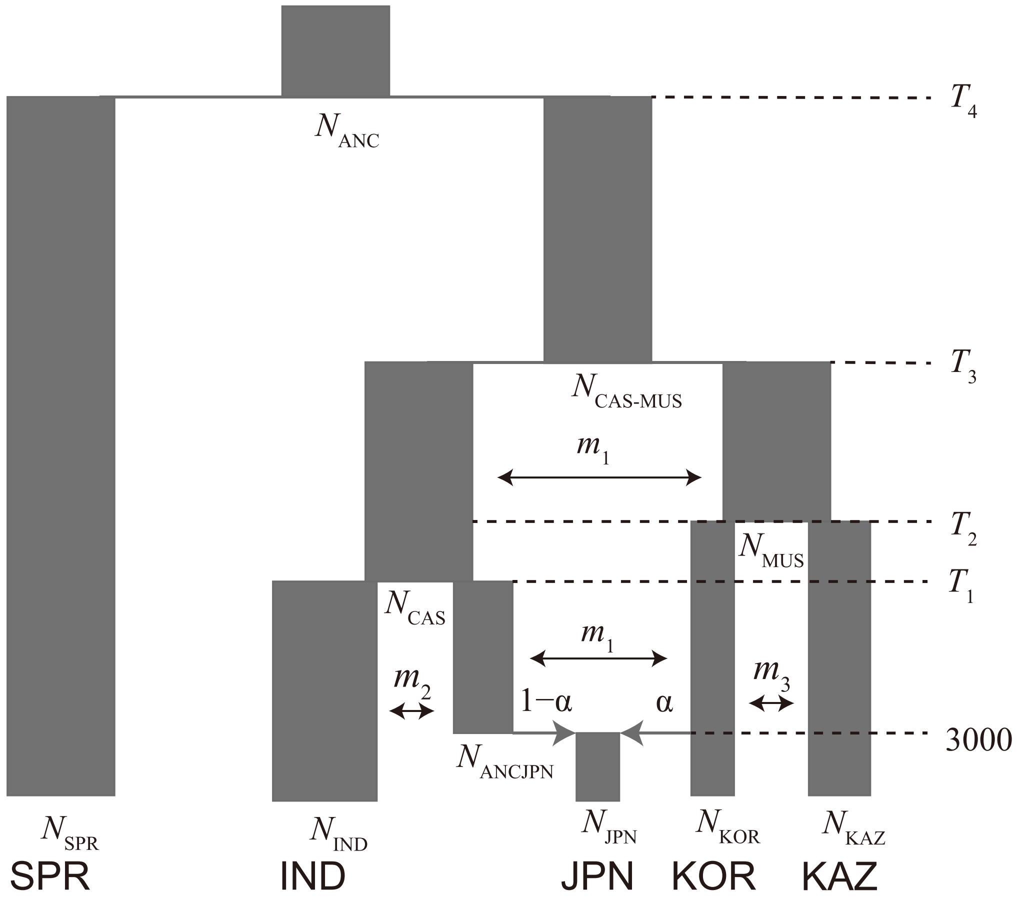


**Supplementary Figure 6.**

The demographic model we used to estimate the population genetic parameters. The parameters were estimated using the site frequency spectrum data of the five populations shown in the figure. See Supplementary Note 2 for more detail.

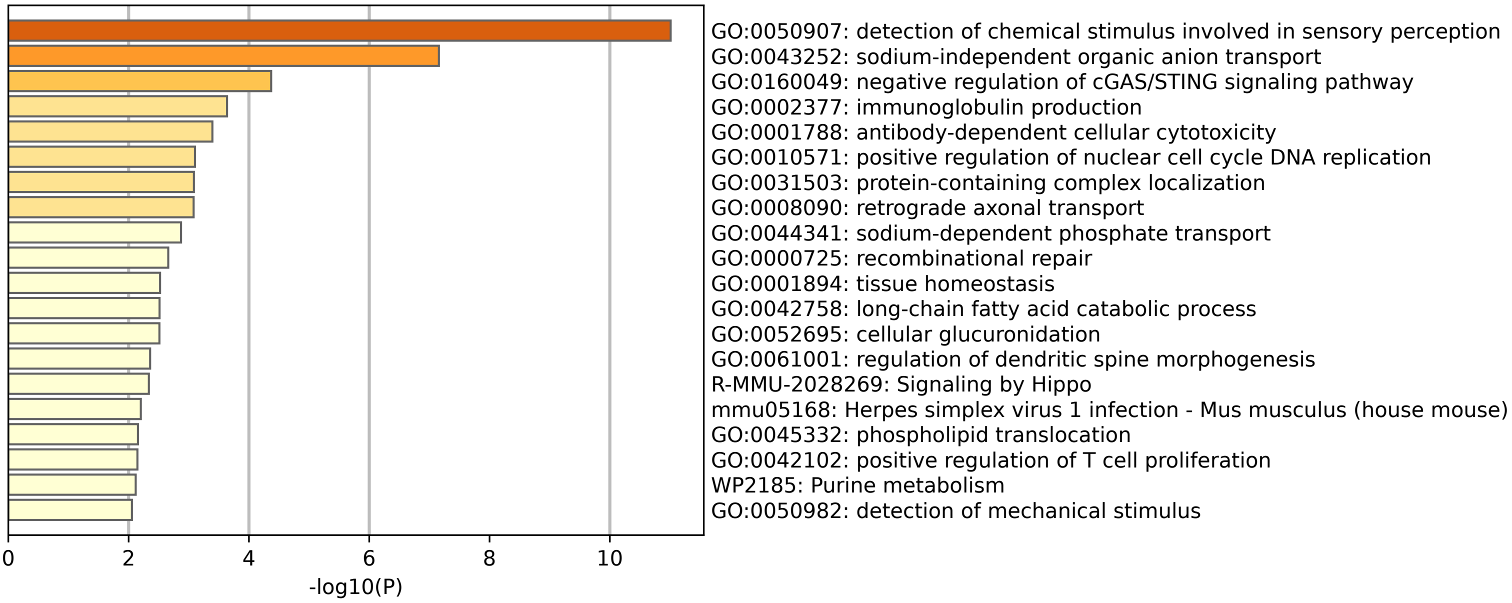


**Supplementary Figure 7.**
Functional enrichment analysis of genes with *castaneus*-ancestry bias, analyzing a total of 842 genes. Gene categories with a *p*-value less than 0.01 are presented.

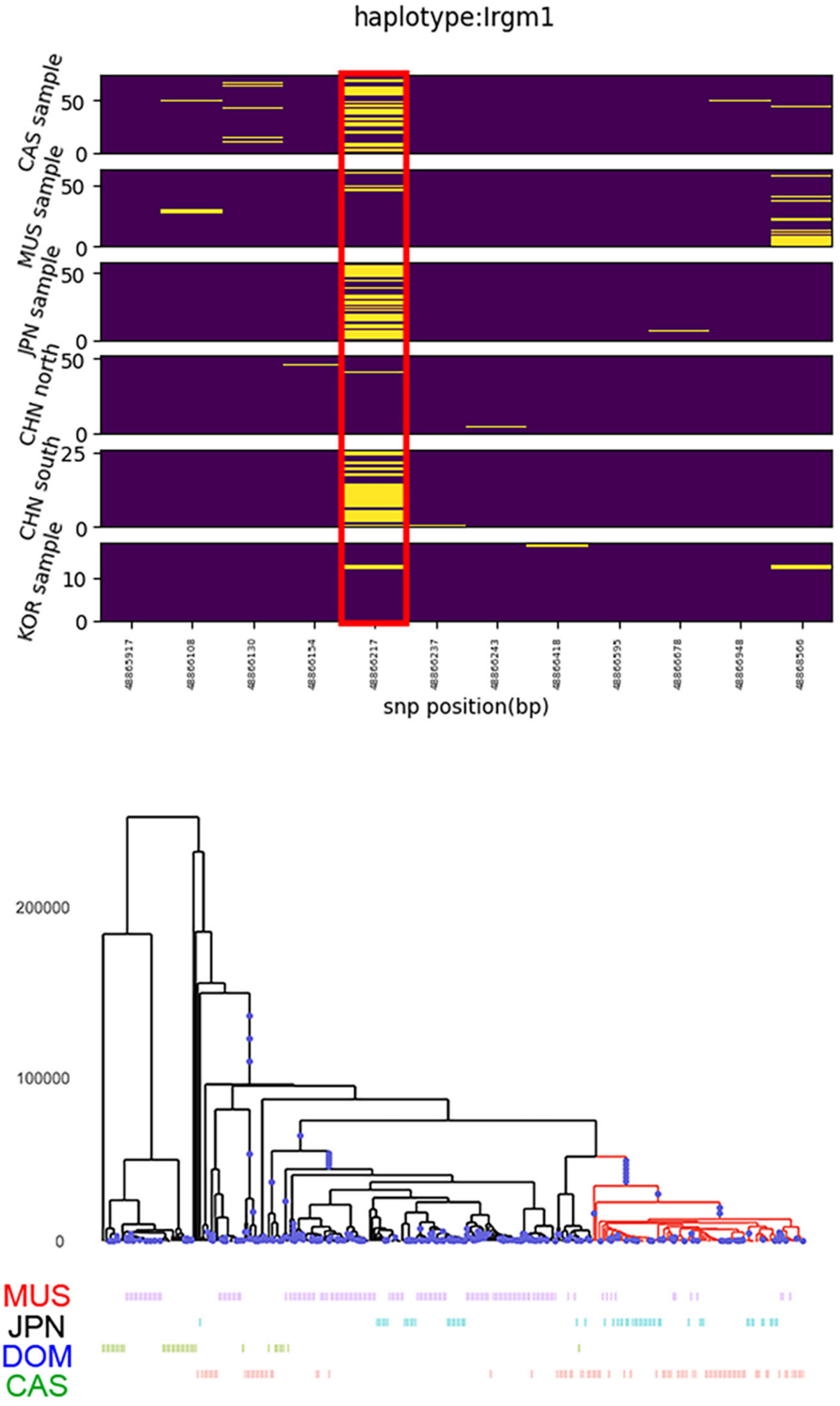


**Supplementary Figure 8.**

Haplotype structure of nonsynonymous SNVs and genealogies of wild house mouse *Irgm1*. The upper panel shows the segregation pattern of nonsynonymous SNVs in *Irgm1*. The site in the red rectangle represents the focal site that showed *castaneus*-ancestry enrichment in the Japanese (JPN) samples. CAS, MUS, KOR and CHN represent *castaneus*-type, *musculus*-type, Korean and Chinese samples, respectively. The lower panel shows the inferred genealogy around the focal site. The branches with the derived mutation are colored in red. The labels of samples are shown at the bottom of the tree. CAS, DOM, MUS, and JPN represent *castaneus*-type, *domesticus*-type, *musculus*-type, and Japanese samples, respectively.

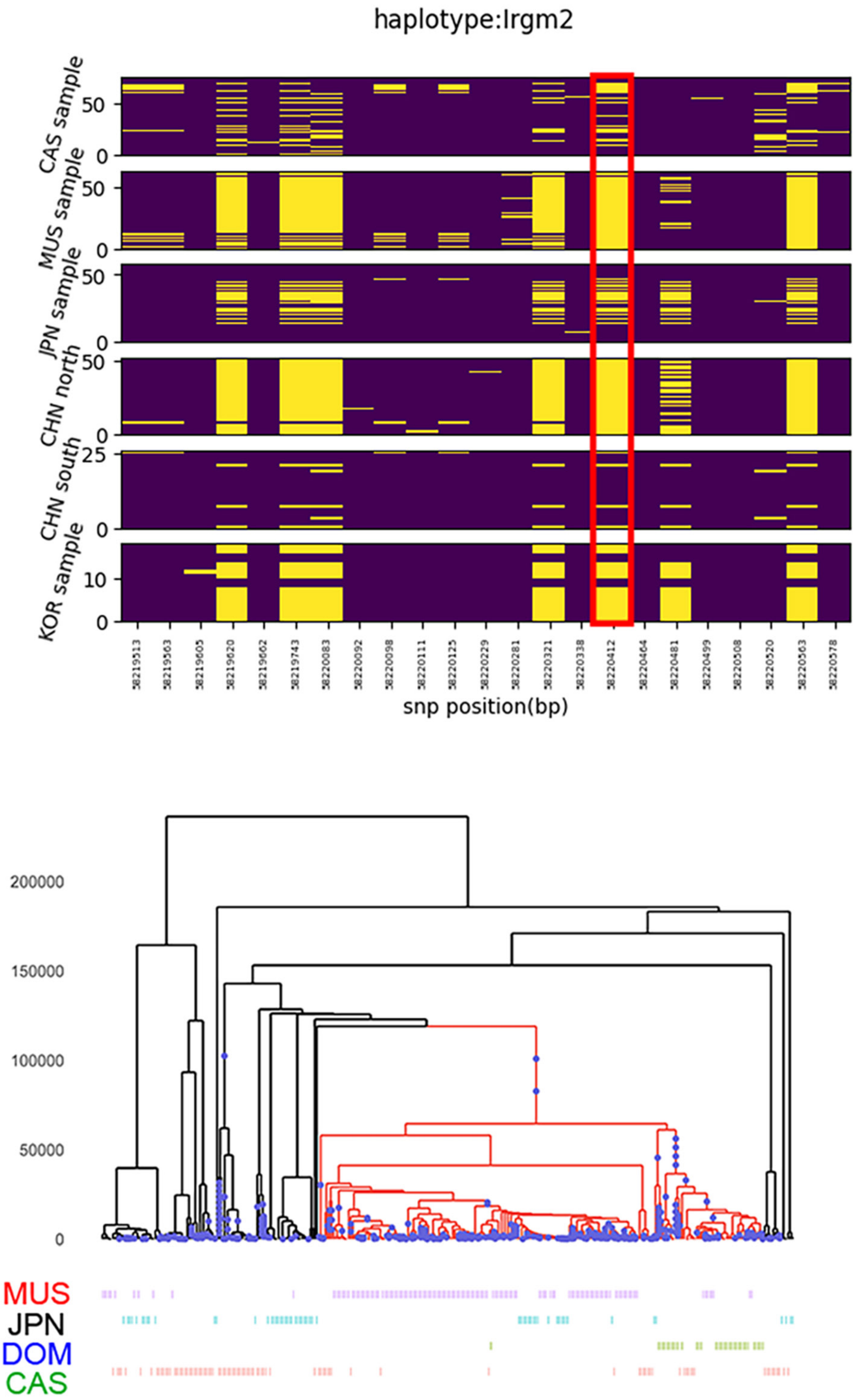


**Supplementary Figure 9.**

Haplotype structure of nonsynonymous SNVs and genealogies of wild house mouse *Irgm2*. The upper panel shows the segregation pattern of nonsynonymous SNVs in *Irgm2*. The site in the red rectangle represents the focal site that showed *castaneus*-ancestry enrichment in the Japanese (JPN) samples. CAS, MUS, KOR and CHN represent *castaneus*-type, *musculus*-type, Korean and Chinese samples, respectively. The lower panel shows the inferred genealogy around the focal site. The branches with the derived mutation are colored in red. The labels of samples are shown at the bottom of the tree. CAS, DOM, MUS, and JPN represent *castaneus*-type, *domesticus*-type, *musculus*-type, and Japanese samples, respectively.

**Supplementary Table 1. List of samples used in this study.**

| Sample ID | Place (country) | Place (city) | Mitochondrial DNA haplogroups | Chromosome Y haplogroups | Species | Sex | Latitude | Longitude | Mean Coverage |
|---|---|---|---|---|---|---|---|---|---|
| HI0111 | Indonesia | Bogor | *castaneus* | | *Mus musculus* | female | -6.6 | 106.8 | 29.0 |
| HI0196 | Philippines | Catarman | *castaneus* | | *Mus musculus* | female | 12.45 | 124.65 | 28.7 |
| HI0410 | South Korea | Chuncheon | *musculus* | *musculus* | *Mus musculus* | male | 37.866667 | 127.733333 | 23.4 |
| HI0447 | Philippines | Languna | *castaneus* | | *Mus musculus* | female | 14.17 | 121.22 | 24.4 |
| HI0468 | South Korea | Chuncheon | *musculus* | *musculus* | *Mus musculus* | male | 37.866667 | 127.733333 | 23.7 |
| HS1413 | Russia | Kraskino | *musculus* | *musculus* | *Mus musculus* | male | 42.711194 | 130.787056 | 25.7 |
| HS3540 | South Korea | Busan | *musculus* | | *Mus musculus* | female | 35.1 | 129.033333 | 30.8 |
| HS3603 | Japan | Kagoshima | *castaneus* | *musculus* | *Mus musculus* | male | 31.596806 | 130.557139 | 29.7 |
| HS3721 | Myanmar | Lashio | *castaneus* | | *Mus musculus* | female | 22.933333 | 97.75 | 30.6 |
| HS5817 | Japan | Fukui | *musculus* | | *Mus musculus* | female | 36.064056 | 136.219583 | 30.6 |
| HS5821 | Japan | Kumamoto | *musculus* | | *Mus musculus* | female | 32.803 | 130.707861 | 27.8 |
| HS5822 | Japan | Yamanashi | *musculus* | | *Mus musculus* | female | 35.693444 | 138.686889 | 26.6 |
| HS6021 | Japan | Saga | *musculus* | *musculus* | *Mus musculus* | male | 33.2635 | 130.300833 | 21.4 |
| HS6022 | Japan | Oshu | *castaneus* | *musculus* | *Mus musculus* | male | 39.144472 | 141.139139 | 24.0 |
| HS6023 | Japan | Sendai | *musculus* | | *Mus musculus* | female | 38.268222 | 140.869417 | 23.3 |
| HS6025 | Japan | Hiroshima | *musculus* | | *Mus musculus* | female | 34.38525 | 132.455306 | 24.7 |
| HS6026 | Japan | Tsukuba | *musculus* | | *Mus musculus* | female | 36.083472 | 140.076444 | 22.1 |
| MG0294 | China | Changchun | *musculus* | | *Mus musculus* | female | 43.897 | 125.326 | 29.0 |
| MG0295 | China | Chengdu | *musculus* | *musculus* | *Mus musculus* | male | 30.66 | 104.063333 | 29.1 |
| MG0439 | China | Shanghai | *castaneus* | *musculus* | *Mus musculus* | male | 31.228611 | 121.474722 | 29.9 |
| MG0488 | Japan | Aizuwakamatsu | *castaneus* | | *Mus musculus* | female | 37.49483 | 139.92975 | 30.1 |
| MG0490 | Japan | Akita | *musculus* | | *Mus musculus* | female | 39.719889 | 140.102583 | 23.4 |
| MG0682 | South Korea | Baengnyeong Island | *musculus* | | *Mus musculus* | female | 37.966667 | 124.65 | 28.7 |
| MG0690 | China | Yinchuan | *musculus* | *musculus* | *Mus musculus* | male | 38.472 | 106.2589 | 24.4 |
| MG0710 | China | Haikou | *castaneus* | | *Mus musculus* | female | 20.042778 | 110.341667 | 25.3 |
| MG0711 | China | Sanya | *castaneus* | | *Mus musculus* | female | 18.2529 | 109.5117 | 24.8 |
| MG0783 | China | Jinan | *musculus* | *musculus* | *Mus musculus* | male | 36.6702 | 117.0207 | 24.6 |
| MG0810 | China | Tongliao | *musculus* | *musculus* | *Mus musculus* | male | 43.654 | 122.243 | 30.0 |
| MG0826 | China | Tianjin | *musculus* | *musculus* | *Mus musculus* | male | 39.123611 | 117.198056 | 24.1 |
| MG0869 | China | Golmud | *musculus* | | *Mus musculus* | female | 36.4072 | 94.9283 | 24.0 |
| MG2021 | China | Tianjin | *musculus* | *musculus* | *Mus musculus* | male | 39.123611 | 117.198056 | 23.9 |
| MG2024 | China | Shijiazhuang | *musculus* | | *Mus musculus* | female | 38.0377 | 114.5304 | 24.0 |
| MG2026 | China | Tumen | *musculus* | | *Mus musculus* | female | 42.966667 | 129.85 | 24.6 |
| MG2030 | China | Xilinhot | *musculus* | | *Mus musculus* | female | 43.9334 | 116.086 | 24.4 |
| MG2031 | China | Laiyang | *musculus* | *musculus* | *Mus musculus* | male | 36.975833 | 120.713611 | 24.6 |
| MG3046 | Russia | South Sakhalin | *castaneus* | | *Mus musculus* | female | 46.966667 | 142.733333 | 29.1 |
| MG3047 | Russia | South Sakhalin | *castaneus* | *musculus* | *Mus musculus* | male | 46.966667 | 142.733333 | 28.9 |
| HI0112 | Indonesia | Bogor | *domesticus* | *castaneus* | *Mus musculus* | male | | Fujiwara et al. 2022a | |
| HI0115 | Indonesia | Denpasar | *castaneus* | | *Mus musculus* | female | | Fujiwara et al. 2022a | |
| HI0116 | Indonesia | Denpasar | *castaneus* | | *Mus musculus* | female | | Fujiwara et al. 2022a | |
| HI0134 | Indonesia | Lembang | *castaneus* | *musculus* | *Mus musculus* | male | | Fujiwara et al. 2022a | |
| HI0159 | India | Leh | *castaneus* | *castaneus* | *Mus musculus* | male | | Fujiwara et al. 2022a | |
| HI0161 | India | Leh | *castaneus* | *castaneus* | *Mus musculus* | male | | Fujiwara et al. 2022a | |
| HI0173 | Pakistan | Islamabad | *castaneus* | | *Mus musculus* | female | | Fujiwara et al. 2022a | |
| HI0175 | Pakistan | Islamabad | *castaneus* | | *Mus musculus* | female | | Fujiwara et al. 2022a | |
| HI0185 | India | Delhi | *castaneus* | *castaneus* | *Mus musculus* | male | | Fujiwara et al. 2022a | |
| HI0261 | Pakistan | Sahiwal | *castaneus* | *castaneus* | *Mus musculus* | male | | Fujiwara et al. 2022a | |
| HI0264 | Pakistan | Lahore | *castaneus* | *castaneus* | *Mus musculus* | male | | Fujiwara et al. 2022a | |
| HI0274 | India | Mysore | *castaneus* | | *Mus musculus* | female | | Fujiwara et al. 2022a | |
| HI0313 | India | Bhubaneswar | *castaneus* | | *Mus musculus* | female | | Fujiwara et al. 2022a | |
| HI0321 | India | Bhubaneswar | *castaneus* | *castaneus* | *Mus musculus* | male | | Fujiwara et al. 2022a | |
| HI0328 | India | Hyderabad | *castaneus* | *castaneus* | *Mus musculus* | male | | Fujiwara et al. 2022a | |
| HI0329 | India | Hyderabad | *castaneus* | | *Mus musculus* | female | | Fujiwara et al. 2022a | |
| HI0347 | Bangladesh | Mymensingh | *castaneus* | | *Mus musculus* | female | | Fujiwara et al. 2022a | |
| HI0373 | Bangladesh | Dhaka | *castaneus* | | *Mus musculus* | female | | Fujiwara et al. 2022a | |
| HI0484 | Sri Lanka | Peradeniya | *castaneus* | *castaneus* | *Mus musculus* | male | | Fujiwara et al. 2022a | |
| HI0488 | Sri Lanka | Colombo | *castaneus* | *castaneus* | *Mus musculus* | male | | Fujiwara et al. 2022a | |
| HI0505 | Vietnam | Vinh Phuc | *castaneus* | *castaneus* | *Mus musculus* | male | | Fujiwara et al. 2022a | |
| HI0520 | Vietnam | Hanoi | *castaneus* | *castaneus* | *Mus musculus* | male | | Fujiwara et al. 2022a | |
| HS0682 | South Korea | Baengnyeong Island | *musculus* | *musculus* | *Mus musculus* | male | | Fujiwara et al. 2022a | |
| HS1368 | South Korea | Gyeongju | *musculus* | *musculus* | *Mus musculus* | male | | Fujiwara et al. 2022a | |
| HS1411 | Russia | Khasan | *castaneus* | *musculus* | *Mus musculus* | male | | Fujiwara et al. 2022a | |
| HS1464 | Kazakhstan | Aktobe | *musculus* | *musculus* | *Mus musculus* | male | | Fujiwara et al. 2022a | |
| HS1467 | Nepal | Tukuche | Nepal | *castaneus* | *Mus musculus* | male | | Fujiwara et al. 2022a | |
| HS1523 | Nepal | Kathmandu | Nepal | *castaneus* | *Mus musculus* | male | | Fujiwara et al. 2022a | |
| HS2323 | Japan | Hakodate | *musculus* | *musculus* | *Mus musculus* | male | | Fujiwara et al. 2022a | |
| HS2326 | Japan | Nayoro | *castaneus* | *musculus* | *Mus musculus* | male | | Fujiwara et al. 2022a | |
| HS2400 | Taiwan | Taitung | *castaneus* | *musculus* | *Mus musculus* | male | | Fujiwara et al. 2022a | |
| HS2445 | Japan | Takikawa | *castaneus* | *musculus* | *Mus musculus* | male | | Fujiwara et al. 2022a | |
| HS2788 | Japan | Shuri | *musculus* | *musculus* | *Mus musculus* | male | | Fujiwara et al. 2022a | |
| HS2815 | Japan | Nonoichi | *musculus* | *musculus* | *Mus musculus* | male | | Fujiwara et al. 2022a | |
| HS3534 | Japan | Yamagata | *musculus* | *musculus* | *Mus musculus* | male | | Fujiwara et al. 2022a | |
| HS3538 | Japan | Tono | *musculus* | | *Mus musculus* | female | | Fujiwara et al. 2022a | |
| HS3604 | Russia | Tomsk | *domesticus* | | *Mus musculus* | female | | Fujiwara et al. 2022a | |
| HS3605 | Russia | Gorno-Altaysk | *musculus* | *musculus* | *Mus musculus* | male | | Fujiwara et al. 2022a | |
| HS3607 | Russia | Okha | *domesticus* | *musculus* | *Mus musculus* | male | | Fujiwara et al. 2022a | |
| HS3608 | Russia | Irkutsk | *musculus* | *musculus* | *Mus musculus* | male | | Fujiwara et al. 2022a | |
| HS3612 | Russia | Astrakhan | *musculus* | *musculus* | *Mus musculus* | male | | Fujiwara et al. 2022a | |
| HS4056 | Japan | Fujisawa | *musculus* | | *Mus musculus* | female | | Fujiwara et al. 2022a | |
| HS4097 | Japan | Misasa | *musculus* | *musculus* | *Mus musculus* | male | | Fujiwara et al. 2022a | |
| HS4120 | Japan | Ishii | *musculus* | | *Mus musculus* | female | | Fujiwara et al. 2022a | |
| HS4169 | Japan | Tsukuba | *musculus* | *musculus* | *Mus musculus* | male | | Fujiwara et al. 2022a | |
| HS4233 | South Korea | Hwacheon | *musculus* | *musculus* | *Mus musculus* | male | | Fujiwara et al. 2022a | |
| HS4238 | South Korea | Ganghwa Island | *musculus* | | *Mus musculus* | female | | Fujiwara et al. 2022a | |
| HS4271 | Japan | Setana | *musculus* | *musculus* | *Mus musculus* | male | | Fujiwara et al. 2022a | |
| HS4272 | Japan | Fukuoka | *musculus* | *musculus* | *Mus musculus* | male | | Fujiwara et al. 2022a | |
| HS4273 | Japan | Yamaguchi | *musculus* | | *Mus musculus* | female | | Fujiwara et al. 2022a | |
| HS4411 | Japan | Oda | *musculus* | *musculus* | *Mus musculus* | male | | Fujiwara et al. 2022a | |
| MG0230 | Japan | Ashiro | *castaneus* | *musculus* | *Mus musculus* | male | | Fujiwara et al. 2022a | |
| MG0296 | Japan | Sapporo | *castaneus* | *musculus* | *Mus musculus* | male | | Fujiwara et al. 2022a | |
| MG0297 | Japan | Sapporo | *castaneus* | | *Mus musculus* | female | | Fujiwara et al. 2022a | |

**Supplementary Table 1. (continued).**

| | | | | | | | |
|---|---|---|---|---|---|---|---|
| MG0417 | Iran | Mashhad | *musculus* | *musculus* | *Mus musculus* | male | Fujiwara et al. 2022a |
| MG0444 | South Korea | Busan | *musculus* | *musculus* | *Mus musculus* | male | Fujiwara et al. 2022a |
| MG0501 | China | Guilin | *castaneus* | | *Mus musculus* | female | Fujiwara et al. 2022a |
| MG0504 | China | Guangzhou | *castaneus* | *musculus* | *Mus musculus* | male | Fujiwara et al. 2022a |
| MG0507 | China | Lanzhou | *musculus* | *musculus* | *Mus musculus* | male | Fujiwara et al. 2022a |
| MG0529 | China | Kunming | *castaneus* | *musculus* | *Mus musculus* | male | Fujiwara et al. 2022a |
| MG0565 | China | Xining | *musculus* | | *Mus musculus* | female | Fujiwara et al. 2022a |
| MG0577 | China | Urumqi | *musculus* | | *Mus musculus* | female | Fujiwara et al. 2022a |
| MG0597 | China | Manas | *musculus* | *musculus* | *Mus musculus* | male | Fujiwara et al. 2022a |
| MG0608 | China | Aksu | *musculus* | *musculus* | *Mus musculus* | male | Fujiwara et al. 2022a |
| MG0611 | China | Tacheng | *musculus* | *musculus* | *Mus musculus* | male | Fujiwara et al. 2022a |
| MG0631 | China | Mohe | *musculus* | | *Mus musculus* | female | Fujiwara et al. 2022a |
| MG0686 | China | Kashgar | *musculus* | *musculus* | *Mus musculus* | male | Fujiwara et al. 2022a |
| MG0709 | China | Chongqing | *castaneus* | | *Mus musculus* | female | Fujiwara et al. 2022a |
| MG0713 | China | Zhenjiang | *castaneus* | *musculus* | *Mus musculus* | male | Fujiwara et al. 2022a |
| MG0716 | China | Lhasa | *musculus* | *musculus* | *Mus musculus* | male | Fujiwara et al. 2022a |
| MG0721 | China | Lhasa | *musculus* | | *Mus musculus* | female | Fujiwara et al. 2022a |
| MG0747 | China | Hotan | *musculus* | | *Mus musculus* | female | Fujiwara et al. 2022a |
| MG0795 | China | Ningbo | *castaneus* | | *Mus musculus* | female | Fujiwara et al. 2022a |
| MG0797 | China | Dali | *castaneus* | *musculus* | *Mus musculus* | male | Fujiwara et al. 2022a |
| MG0863 | China | Manzhouli | *castaneus* | *musculus* | *Mus musculus* | male | Fujiwara et al. 2022a |
| MG0871 | China | Dunhuang | *musculus* | *musculus* | *Mus musculus* | male | Fujiwara et al. 2022a |
| MG0908 | China | Wuhan | *castaneus* | *musculus* | *Mus musculus* | male | Fujiwara et al. 2022a |
| MG0917 | China | Lijiang | *castaneus* | *musculus* | *Mus musculus* | male | Fujiwara et al. 2022a |
| MG0992 | China | Qiqihar | *musculus* | | *Mus musculus* | female | Fujiwara et al. 2022a |
| MG3007 | Russia | Krementui | *musculus* | *musculus* | *Mus musculus* | male | Fujiwara et al. 2022a |
| MG3008 | Russia | Krementui | *musculus* | | *Mus musculus* | female | Fujiwara et al. 2022a |
| MG3010 | Russia | Nouth Caucastaneusus | *musculus* | *musculus* | *Mus musculus* | male | Fujiwara et al. 2022a |
| MG3012 | Russia | Chita | *musculus* | | *Mus musculus* | female | Fujiwara et al. 2022a |
| MG3018 | Russia | Rudnaya Pristan | *castaneus* | *musculus* | *Mus musculus* | male | Fujiwara et al. 2022a |
| MG3025 | Russia | Vladivostok | *castaneus* | | *Mus musculus* | female | Fujiwara et al. 2022a |
| MG3026 | Russia | Birakan | *musculus* | *musculus* | *Mus musculus* | male | Fujiwara et al. 2022a |
| MG3037 | Russia | Khabarovsk | *domesticus* | | *Mus musculus* | female | Fujiwara et al. 2022a |
| MG3044 | Estonia | Tallinn | *musculus* | *musculus* | *Mus musculus* | male | Fujiwara et al. 2022a |
| MG3045 | Russia | Poronaysk | *musculus* | | *Mus musculus* | female | Fujiwara et al. 2022a |
| MG3048 | Russia | Yuzhno-Sakhalinsk | *musculus* | | *Mus musculus* | female | Fujiwara et al. 2022a |
| MG3056 | Russia | Moscow | *domesticus* | | *Mus musculus* | female | Fujiwara et al. 2022a |
| MG3066 | Ukraine | Donetsk | *musculus* | *musculus* | *Mus musculus* | male | Fujiwara et al. 2022a |
| MG5086 | China | Jiayuguan | *musculus* | *musculus* | *Mus musculus* | male | Fujiwara et al. 2022a |
| MG5135 | Iran | Now Shahr | *castaneus* | | *Mus musculus* | female | Fujiwara et al. 2022a |
| HS5555 | Madagascar | Antanànarivo | Madagascar | *castaneus* | *Mus musculus* | male | Fujiwara et al. 2022b |
| HS5558 | Madagascar | Antanànarivo | Madagascar | | *Mus musculus* | female | Fujiwara et al. 2022b |
| HS5559 | Madagascar | Antanànarivo | Madagascar | *castaneus* | *Mus musculus* | male | Fujiwara et al. 2022b |
| HS5560 | Madagascar | Antanànarivo | Madagascar | *castaneus* | *Mus musculus* | male | Fujiwara et al. 2022b |
| HS5561 | Madagascar | Antanànarivo | Madagascar | | *Mus musculus* | female | Fujiwara et al. 2022b |
| ERS003044 | India | Himalaya | *castaneus* | | *Mus musculus* | female | Harr et al. 2016 |
| ERS739373 | Germany | Cologne-Bonn | *domesticus* | *domesticus* | *Mus musculus* | male | Harr et al. 2016 |
| ERS739374 | Germany | Cologne-Bonn | *domesticus* | *domesticus* | *Mus musculus* | male | Harr et al. 2016 |
| ERS739375 | Germany | Cologne-Bonn | *domesticus* | *domesticus* | *Mus musculus* | male | Harr et al. 2016 |
| ERS739376 | Germany | Cologne-Bonn | *domesticus* | *domesticus* | *Mus musculus* | male | Harr et al. 2016 |
| ERS739377 | Germany | Cologne-Bonn | *domesticus* | *domesticus* | *Mus musculus* | male | Harr et al. 2016 |
| ERS739378 | Germany | Cologne-Bonn | *domesticus* | *domesticus* | *Mus musculus* | male | Harr et al. 2016 |
| ERS739379 | Germany | Cologne-Bonn | *domesticus* | *domesticus* | *Mus musculus* | male | Harr et al. 2016 |
| ERS739382 | France | Massif Central | *domesticus* | *domesticus* | *Mus musculus* | male | Harr et al. 2016 |
| ERS739385 | France | Massif Central | *domesticus* | *domesticus* | *Mus musculus* | male | Harr et al. 2016 |
| ERS739386 | France | Massif Central | *domesticus* | *domesticus* | *Mus musculus* | male | Harr et al. 2016 |
| ERS739388 | France | Massif Central | *domesticus* | *domesticus* | *Mus musculus* | male | Harr et al. 2016 |
| ERS739389 | Iran | Ahvaz | *domesticus* | *domesticus* | *Mus musculus* | male | Harr et al. 2016 |
| ERS739393 | Iran | Ahvaz | *domesticus* | *domesticus* | *Mus musculus* | male | Harr et al. 2016 |
| ERS739395 | Iran | Ahvaz | *domesticus* | *domesticus* | *Mus musculus* | male | Harr et al. 2016 |
| ERS739396 | Iran | Ahvaz | *domesticus* | *domesticus* | *Mus musculus* | male | Harr et al. 2016 |
| ERS957497 | Czech Republic | Studenec | *musculus* | | *Mus musculus* | female | Harr et al. 2016 |
| ERS957498 | Czech Republic | Studenec | *musculus* | *musculus* | *Mus musculus* | male | Harr et al. 2016 |
| ERS957499 | Czech Republic | Studenec | *musculus* | | *Mus musculus* | female | Harr et al. 2016 |
| ERS957500 | Czech Republic | Studenec | *musculus* | | *Mus musculus* | female | Harr et al. 2016 |
| ERS957501 | Czech Republic | Studenec | *musculus* | | *Mus musculus* | female | Harr et al. 2016 |
| ERS957502 | Czech Republic | Studenec | *musculus* | *musculus* | *Mus musculus* | male | Harr et al. 2016 |
| ERS957503 | Czech Republic | Studenec | *musculus* | | *Mus musculus* | female | Harr et al. 2016 |
| ERS957504 | Kazakhstan | Almaty | *musculus* | | *Mus musculus* | female | Harr et al. 2016 |
| ERS957506 | Kazakhstan | Almaty | *musculus* | | *Mus musculus* | female | Harr et al. 2016 |
| ERS957508 | Kazakhstan | Almaty | *musculus* | *musculus* | *Mus musculus* | male | Harr et al. 2016 |
| ERS957510 | Kazakhstan | Almaty | *musculus* | *musculus* | *Mus musculus* | male | Harr et al. 2016 |
| ERS957512 | Spain | Madrid | *domesticus* | *Mus spretus* | *Mus spretus* | male | Harr et al. 2016 |
| ERS957513 | Spain | Madrid | *Mus spretus* | *Mus spretus* | *Mus spretus* | male | Harr et al. 2016 |
| ERS957514 | Spain | Madrid | *Mus spretus* | *Mus spretus* | *Mus spretus* | male | Harr et al. 2016 |
| ERS957515 | Spain | Madrid | *Mus spretus* | | *Mus spretus* | female | Harr et al. 2016 |
| ERS957516 | Spain | Madrid | *Mus spretus* | | *Mus spretus* | female | Harr et al. 2016 |
| ERS957517 | Spain | Madrid | *Mus spretus* | *Mus spretus* | *Mus spretus* | male | Harr et al. 2016 |
| ERS957519 | Spain | Madrid | *Mus spretus* | *Mus spretus* | *Mus spretus* | male | Harr et al. 2016 |

Fujiwara, K., Kawai, Y., Takada, T. et al. Insights into *Mus musculus* Population Structure across Eurasia Revealed by Whole-Genome Analysis. Genome Biol. Evol. **14**, evac068 (2022a)
Fujiwara, K., Ranorosoa, M. C., Ohdachi, S. D. et al. Whole-genome sequencing analysis of wild house mice (*Mus musculus*) captured in Madagascar. Genes Genet. Syst. **97**, 193–207 (2022b)
Harr, B., Karakoc, E., Neme, R. et al. Genomic resources for wild populations of the house mouse, *Mus musculus* and its close relative *Mus spretus*. Sci Data **3**, 160075 (2016).

**Supplementray Table 2. Allele frequency of introgressed alleles with different *r***

| $\alpha$ | $\beta$ | $\gamma$ | $F_X$ | $F_Y$ | $F_M$ |
|---|---|---|---|---|---|
| 0 | 0 | 0 | 0.999 | 1 | 0.005 |
| 1 | 0 | 0 | 0.851 | 1 | 0 |
| 0 | 1 | 0 | 0.645 | 1 | 0.006 |
| 0 | 0 | 1 | 0.975 | 1 | 0 |
| 1 | 1 | 0 | 0.146 | 0.995 | 0.001 |
| 1 | 0 | 1 | 0.447 | 1 | 0 |
| 0 | 1 | 1 | 0.298 | 0.996 | 0 |

*$F_X$, $F_Y$, and $F_M$ represent the average allele frequency of introgressed X-chromosomal, Y-chromosomal, and mitochondrial genomes after $0.1N$ generations, respectively ($\alpha = 0, \beta = 1, \gamma = 0$).

**Supplementary Table 3. Estimated population genetic parameters and confidence intervals**

| Parameter | MLE | 95% CI (lower bound) | 95% CI (upper bound) |
|---|---|---|---|
| $N_{\mathrm{SPR}}$ | 264,382 | 268,833 | 301,671 |
| $N_{\mathrm{IND}}$ | 180,130 | 221,839 | 376,998 |
| $N_{\mathrm{ANCJPN}}$ | 288,068 | 870,163 | 997,358 |
| $N_{\mathrm{JPN}}$ | 33,517 | 36,201 | 43,226 |
| $N_{\mathrm{KOR}}$ | 38,806 | 42,835 | 54,087 |
| $N_{\mathrm{KAZ}}$ | 98,772 | 125,476 | 150,902 |
| $N_{\mathrm{CAS}}$ | 82,293 | 406,286 | 886,234 |
| $N_{\mathrm{MUS}}$ | 67,719 | 85,905 | 139,314 |
| $N_{\mathrm{CAS\text{-}MUS}}$ | 577,379 | 587,568 | 728,170 |
| $N_{\mathrm{ANC}}$ | 789,873 | 939,508 | 939,508 |
| $T_1$ | 41,315 | 65,511 | 89,629 |
| $T_2$ | 166,065 | 140,655 | 187,281 |
| $T_3$ | 428,735 | 149,086 | 394,045 |
| $T_4$ | 1,779,763 | 693,233 | 1,988,374 |
| $m_1$ | $9.73\times10^{-9}$ | $6.18\times10^{-10}$ | $1.15\times10^{-3}$ |
| $m_2$ | $9.91\times10^{-6}$ | $8.79\times10^{-6}$ | $1.04\times10^{-5}$ |
| $m_3$ | $9.40\times10^{-10}$ | $4.08\times10^{-6}$ | $1.19\times10^{-6}$ |
| $\alpha$ | 0.864 | 0.882 | 0.887 |

**Supplementary Table 4. List of genes in the *castaneus*-enriched 20-kb-lengh windows.**

| Gene symbol | Chromosome* | Exon start | Exon end | *α* ** | *p*-value |
|---|---|---|---|---|---|
| Igkv14-130 | chr6 | 67791216 | 67791511 | 0.00034143 | <0.001 |
| Ano4 | chr10 | 89116865 | 89117002 | 0.000408037 | <0.001 |
| Kcnab1 | chr3 | 65302147 | 65302192 | 0.00228414 | <0.001 |
| Ip6k3 | chr17 | 27144881 | 27145322 | 0.003331868 | <0.001 |
| Tmprss15 | chr16 | 79087448 | 79087516 | 0.009323779 | 0.001 |
| 4930430A15Rik | chr2 | 111203949 | 111203977 | 0.015082046 | 0.001 |
| Npffr2 | chr5 | 89567815 | 89568143 | 0.016707939 | 0.001 |
| Olfr1135 | chr2 | 87671432 | 87672365 | 0.017028194 | 0.001 |
| Gm19965 | chr1 | 116820828 | 116822762 | 0.018031436 | 0.001 |
| Olfr290 | chr7 | 84915780 | 84916728 | 0.021504683 | 0.001 |
| Rnaseh2b | chr14 | 62350399 | 62350476 | 0.024011899 | 0.001 |
| Igkv11-125 | chr6 | 67913754 | 67914052 | 0.026437782 | 0.001 |
| Slco1b2 | chr6 | 141676203 | 141676385 | 0.026629043 | 0.001 |
| Utp4 | chr8 | 106898059 | 106898251 | 0.028524644 | 0.002 |
| Vmn2r70 | chr7 | 85565999 | 85566127 | 0.031127151 | 0.002 |
| Igkv4-91 | chr6 | 68768554 | 68768863 | 0.03188334 | 0.002 |
| Igkv4-92 | chr6 | 68755544 | 68755593 | 0.03188334 | 0.002 |
| Bank1 | chr3 | 136064112 | 136064176 | 0.037243547 | 0.002 |
| Dglucy | chr12 | 100829628 | 100829725 | 0.037397923 | 0.002 |
| Sh3bp5l | chr11 | 58345929 | 58346397 | 0.038044341 | 0.002 |
| Olfr130 | chr17 | 38067172 | 38068126 | 0.038698078 | 0.002 |
| Galnt10 | chr11 | 57783689 | 57783839 | 0.038716065 | 0.002 |
| Vmn1r71 | chr7 | 10747835 | 10748759 | 0.03975009 | 0.002 |
| Zfp329 | chr7 | 12807911 | 12807973 | 0.041505866 | 0.002 |
| Col4a4 | chr1 | 82541169 | 82541304 | 0.043248632 | 0.002 |
| Igkv12-41 | chr6 | 69858419 | 69858717 | 0.046921486 | 0.003 |
| Traj50 | chr14 | 54167589 | 54167652 | 0.048255059 | 0.003 |
| Traj59 | chr14 | 54156132 | 54156194 | 0.048255059 | 0.003 |
| Traj58 | chr14 | 54157279 | 54157342 | 0.048255059 | 0.003 |
| Traj57 | chr14 | 54158506 | 54158569 | 0.048255059 | 0.003 |
| Traj56 | chr14 | 54159262 | 54159325 | 0.048255059 | 0.003 |
| Traj53 | chr14 | 54162643 | 54162709 | 0.048255059 | 0.003 |
| Traj52 | chr14 | 54165315 | 54165381 | 0.048255059 | 0.003 |
| Traj47 | chr14 | 54171799 | 54171856 | 0.048255059 | 0.003 |
| Traj48 | chr14 | 54169807 | 54169868 | 0.048255059 | 0.003 |
| Traj46 | chr14 | 54172335 | 54172398 | 0.048255059 | 0.003 |
| Traj44 | chr14 | 54173689 | 54173750 | 0.048255059 | 0.003 |
| Traj43 | chr14 | 54174741 | 54174798 | 0.048255059 | 0.003 |

**Supplementary Table 4. (continued).**

| | | | | | |
|---|---|---|---|---|---|
| Traj42 | chr14 | 54175772 | 54175836 | 0.048255059 | 0.003 |
| Traj49 | chr14 | 54168685 | 54168747 | 0.048255059 | 0.003 |
| Traj45 | chr14 | 54172827 | 54172890 | 0.048255059 | 0.003 |
| Myo1e | chr9 | 70383803 | 70384005 | 0.049179082 | 0.003 |
| Csmd3 | chr15 | 48621993 | 48622106 | 0.053544563 | 0.003 |
| Igkv1-88 | chr6 | 68862264 | 68862577 | 0.055939402 | 0.003 |
| Zfand6 | chr7 | 84634238 | 84634392 | 0.057985751 | 0.003 |
| Olfr1093 | chr2 | 86785731 | 86786700 | 0.060360446 | 0.004 |
| Grm4 | chr17 | 27502646 | 27503165 | 0.062581687 | 0.004 |
| Tspan18 | chr2 | 93211817 | 93211892 | 0.062860807 | 0.004 |
| Olfr510 | chr7 | 108667417 | 108668362 | 0.06655147 | 0.004 |
| Pyurf | chr6 | 57689736 | 57689878 | 0.069202991 | 0.004 |
| Ankfy1 | chr11 | 72712156 | 72712349 | 0.070100183 | 0.004 |
| Olfr1265 | chr2 | 90036920 | 90037850 | 0.07055247 | 0.004 |
| Grik3 | chr4 | 125659688 | 125659802 | 0.070562673 | 0.004 |
| Rxylt1 | chr10 | 122088878 | 122089049 | 0.072692812 | 0.004 |
| Ahrr | chr13 | 74277627 | 74277809 | 0.073008635 | 0.004 |
| Pdlim4 | chr11 | 54056142 | 54056324 | 0.075951724 | 0.005 |
| Slc15a2 | chr16 | 36782271 | 36782359 | 0.079847718 | 0.005 |
| Eaf2 | chr16 | 36793879 | 36793926 | 0.079847718 | 0.005 |
| Scn1a | chr2 | 66280695 | 66280800 | 0.081794982 | 0.005 |
| Sem1 | chr6 | 6558448 | 6558491 | 0.082582822 | 0.005 |
| Clca3a2 | chr3 | 144805675 | 144805940 | 0.082813185 | 0.005 |
| Exoc3 | chr13 | 74206906 | 74207080 | 0.084575273 | 0.005 |
| Mre11a | chr9 | 14832640 | 14832716 | 0.089472476 | 0.006 |
| Fv1 | chr4 | 147868978 | 147870358 | 0.092625995 | 0.006 |
| Miip | chr4 | 147865654 | 147866014 | 0.092625995 | 0.006 |
| Hand1 | chr11 | 57829598 | 57829706 | 0.095460487 | 0.006 |
| Irgm2 | chr11 | 58219484 | 58220672 | 0.096447316 | 0.006 |
| Zfat | chr15 | 68084321 | 68084561 | 0.097728917 | 0.006 |
| Lmna | chr3 | 88484046 | 88484154 | 0.098116647 | 0.006 |
| Cacna2d3 | chr14 | 29467896 | 29467956 | 0.10050976 | 0.006 |
| Cyp4f14 | chr17 | 32914537 | 32914682 | 0.100972194 | 0.006 |
| Chrna9 | chr5 | 65967781 | 65967927 | 0.101281489 | 0.006 |
| Vcan | chr13 | 89731458 | 89731528 | 0.101883435 | 0.006 |
| Ipp | chr4 | 116529670 | 116529808 | 0.10320108 | 0.006 |
| Gm6614 | chr6 | 142011354 | 142011414 | 0.10325781 | 0.006 |
| Tgm6 | chr2 | 130143371 | 130143614 | 0.105243419 | 0.006 |
| Vmn2r1 | chr3 | 64086470 | 64086762 | 0.109977668 | 0.007 |

**Supplementary Table 4. (continued).**

| | | | | | |
|---|---|---|---|---|---|
| Cntnap5b | chr1 | 100358653 | 100358771 | 0.113678213 | 0.007 |
| Tmem41b | chr7 | 109978709 | 109978814 | 0.115914045 | 0.007 |
| Vmn2r76 | chr7 | 86211712 | 86211765 | 0.116040266 | 0.007 |
| Igkv8-34 | chr6 | 70044629 | 70044678 | 0.117030638 | 0.007 |
| Cntln | chr4 | 85033787 | 85033888 | 0.118628902 | 0.007 |
| Spopl | chr2 | 23543196 | 23543324 | 0.11982375 | 0.007 |
| Chd2 | chr7 | 73463635 | 73463775 | 0.119837458 | 0.007 |
| Haus8 | chr8 | 71272283 | 71272312 | 0.123188181 | 0.008 |
| Adcy5 | chr16 | 35274424 | 35274565 | 0.123658634 | 0.008 |
| Ivl | chr3 | 92571349 | 92572756 | 0.125741946 | 0.008 |
| Maea | chr5 | 33365721 | 33365798 | 0.12651257 | 0.008 |
| Uvssa | chr5 | 33379227 | 33379325 | 0.12651257 | 0.008 |
| Olfr630 | chr7 | 103754623 | 103755583 | 0.127386307 | 0.008 |
| Cd59b | chr2 | 104080996 | 104081098 | 0.129823809 | 0.008 |
| Angpt1 | chr15 | 42676164 | 42676461 | 0.130473352 | 0.008 |
| Esco1 | chr18 | 10584247 | 10584362 | 0.131675775 | 0.008 |
| Timm17a | chr1 | 135309134 | 135309198 | 0.132679986 | 0.008 |
| Sap30l | chr11 | 57808023 | 57808122 | 0.135276106 | 0.008 |
| Acadl | chr1 | 66857398 | 66857554 | 0.135425404 | 0.008 |
| Olfr411 | chr11 | 74346634 | 74347582 | 0.135611162 | 0.009 |
| Pkd2 | chr5 | 104503451 | 104503688 | 0.137606236 | 0.009 |
| BC005561 | chr5 | 104517613 | 104522383 | 0.137606236 | 0.009 |
| Slc35c1 | chr2 | 92454214 | 92454774 | 0.13762559 | 0.009 |
| Olfr501-ps1 | chr7 | 108508057 | 108509000 | 0.139897011 | 0.009 |
| Olfr502 | chr7 | 108523003 | 108523948 | 0.139897011 | 0.009 |
| Ilrun | chr17 | 27820176 | 27820334 | 0.142491793 | 0.009 |
| Ttpal | chr2 | 163607226 | 163607291 | 0.144688051 | 0.009 |
| Ino80 | chr2 | 119423604 | 119423776 | 0.144748163 | 0.009 |
| Slc20a1 | chr2 | 129210683 | 129210845 | 0.149280842 | 0.009 |
| Cts7 | chr13 | 61356910 | 61357036 | 0.14960569 | 0.010 |
| Zfp472 | chr17 | 32975913 | 32976040 | 0.150715544 | 0.010 |
| Olfr320 | chr11 | 58683874 | 58684795 | 0.152486374 | 0.010 |
| Olfr29-ps1 | chr4 | 43781365 | 43782328 | 0.15372249 | 0.010 |
| Olfr159 | chr4 | 43770049 | 43771009 | 0.15372249 | 0.010 |
| Gm17067 | chr7 | 42726949 | 42726952 | 0.154499327 | 0.010 |
| Arhgap24 | chr5 | 102552145 | 102552319 | 0.159554778 | 0.010 |
| Igkv12-44 | chr6 | 69815051 | 69815100 | 0.161242273 | 0.010 |
| Igkv5-43 | chr6 | 69823862 | 69823911 | 0.161242273 | 0.010 |
| Cyp4f39 | chr17 | 32468461 | 32468683 | 0.163993683 | 0.010 |

**Supplementary Table 4. (continued).**

| | | | | | |
|---|---|---|---|---|---|
| Tmem126b | chr7 | 90468977 | 90469161 | 0.16567198 | 0.011 |
| Dlg2 | chr7 | 90477856 | 90477875 | 0.16567198 | 0.011 |
| Capn3 | chr2 | 120488073 | 120488157 | 0.165862736 | 0.011 |
| Fam114a2 | chr11 | 57484017 | 57484140 | 0.167016888 | 0.011 |
| Olfr181 | chr16 | 58926184 | 58926569 | 0.16908769 | 0.011 |
| Fam135b | chr15 | 71478186 | 71478260 | 0.170290248 | 0.011 |
| Vmn2r65 | chr7 | 84943140 | 84943264 | 0.171219811 | 0.011 |
| Zc3h6 | chr2 | 128997604 | 128997872 | 0.174238939 | 0.011 |
| Gcfc2 | chr6 | 81942928 | 81943010 | 0.174898186 | 0.011 |
| Olfr299 | chr7 | 86465412 | 86466405 | 0.175379766 | 0.012 |
| Cd209e | chr8 | 3849083 | 3849237 | 0.176848383 | 0.012 |
| Ctnna3 | chr10 | 65002545 | 65002833 | 0.179972387 | 0.012 |
| Cdh9 | chr15 | 16778100 | 16778325 | 0.180892227 | 0.012 |
| Hsf2bp | chr17 | 32011175 | 32011308 | 0.181060632 | 0.012 |
| Zfp455 | chr13 | 67199210 | 67199244 | 0.18170711 | 0.012 |
| Rsl1 | chr13 | 67181748 | 67182947 | 0.18170711 | 0.012 |
| Itgbl1 | chr14 | 123840562 | 123840685 | 0.182941045 | 0.012 |
| Nebl | chr2 | 17590092 | 17590177 | 0.183522088 | 0.012 |
| Vwc2 | chr11 | 11154162 | 11154292 | 0.183611995 | 0.012 |
| Ptprk | chr10 | 28206133 | 28206256 | 0.184162316 | 0.012 |
| Lpcat3 | chr6 | 124700006 | 124700100 | 0.186194798 | 0.012 |
| Emg1 | chr6 | 124704544 | 124704658 | 0.186194798 | 0.012 |
| Bmt2 | chr6 | 13630556 | 13630704 | 0.187402512 | 0.012 |
| Snd1 | chr6 | 28626099 | 28626189 | 0.187573308 | 0.012 |
| Cyp21a1 | chr17 | 34803159 | 34803261 | 0.18773162 | 0.012 |
| Ctsr | chr13 | 61162441 | 61162591 | 0.188172924 | 0.012 |
| Zfp386 | chr12 | 116047857 | 116047866 | 0.188292301 | 0.012 |
| Gdf9 | chr11 | 53436615 | 53437544 | 0.189428272 | 0.013 |
| Leap2 | chr11 | 53422412 | 53422449 | 0.189428272 | 0.013 |
| Uqcrq | chr11 | 53429055 | 53429150 | 0.189428272 | 0.013 |
| Ugt1a9 | chr1 | 88070829 | 88071678 | 0.189822225 | 0.013 |
| Bcl2l15 | chr3 | 103833355 | 103833477 | 0.190418065 | 0.013 |
| Ap4b1 | chr3 | 103821335 | 103821760 | 0.190418065 | 0.013 |
| Gm43064 | chr3 | 103820677 | 103820959 | 0.190418065 | 0.013 |
| Macf1 | chr4 | 123544741 | 123544831 | 0.19637896 | 0.013 |
| Olfr1198 | chr2 | 88745959 | 88746886 | 0.196925345 | 0.013 |
| Igkv4-54 | chr6 | 69632129 | 69632178 | 0.198151271 | 0.013 |
| Igkv4-53 | chr6 | 69648823 | 69649115 | 0.198151271 | 0.013 |
| Vmn2r78 | chr7 | 86915348 | 86915545 | 0.198475984 | 0.013 |

**Supplementary Table 4. (continued).**

| | | | | | |
|---|---|---|---|---|---|
| L3mbtl1 | chr2 | 162971165 | 162971228 | 0.198824038 | 0.013 |
| Ankib1 | chr5 | 3755593 | 3755776 | 0.199533971 | 0.013 |
| Cd55b | chr1 | 130422114 | 130422300 | 0.200829818 | 0.014 |
| Rin2 | chr2 | 145822609 | 145822682 | 0.200873815 | 0.014 |
| Zfp871 | chr17 | 32788137 | 32788140 | 0.201130949 | 0.014 |
| Zfp811 | chr17 | 32797405 | 32798873 | 0.201130949 | 0.014 |
| Me3 | chr7 | 89763319 | 89763353 | 0.20156393 | 0.014 |
| Zfp322a | chr13 | 23357192 | 23357570 | 0.202040359 | 0.014 |
| Nbea | chr3 | 55985609 | 55985778 | 0.202455955 | 0.014 |
| Bsn | chr9 | 108190043 | 108190258 | 0.20278525 | 0.014 |
| Ccdc107 | chr4 | 43495341 | 43495432 | 0.204872389 | 0.014 |
| Car9 | chr4 | 43509117 | 43509210 | 0.204872389 | 0.014 |
| Arhgef39 | chr4 | 43497958 | 43498029 | 0.204872389 | 0.014 |
| Csmd2 | chr4 | 128058961 | 128059178 | 0.205080626 | 0.014 |
| Hlcs | chr16 | 94287791 | 94287926 | 0.205254945 | 0.014 |
| Cyp4f13 | chr17 | 32946804 | 32947002 | 0.205361144 | 0.014 |
| Robo3 | chr9 | 37432841 | 37433001 | 0.205456768 | 0.014 |
| Usp40 | chr1 | 87962423 | 87962488 | 0.205765654 | 0.014 |
| Bend5 | chr4 | 111415174 | 111415400 | 0.205884975 | 0.014 |
| Spag9 | chr11 | 94068935 | 94069035 | 0.207657591 | 0.014 |
| Ccn5 | chr2 | 163832216 | 163832437 | 0.207703416 | 0.014 |
| Tbx20 | chr9 | 24740696 | 24740703 | 0.208159723 | 0.014 |
| Olfr1131 | chr2 | 87628464 | 87629394 | 0.208228103 | 0.014 |
| Cd72 | chr4 | 43454241 | 43454349 | 0.208286224 | 0.014 |
| Ighv7-1 | chr12 | 113896900 | 113896946 | 0.211229464 | 0.015 |
| Igkv5-45 | chr6 | 69776257 | 69776306 | 0.212176404 | 0.015 |
| Asb18 | chr1 | 89968212 | 89968720 | 0.215805063 | 0.015 |
| Fah | chr7 | 84600776 | 84600826 | 0.215882011 | 0.015 |
| Mlh3 | chr12 | 85247611 | 85247723 | 0.215924464 | 0.015 |
| Pdlim5 | chr3 | 142259129 | 142259292 | 0.21673797 | 0.015 |
| Dnah8 | chr17 | 30830772 | 30830889 | 0.217769127 | 0.015 |
| Cnot1 | chr8 | 95788629 | 95788731 | 0.219914566 | 0.016 |
| Olfr1249 | chr2 | 89629939 | 89630896 | 0.220042098 | 0.016 |
| Olfr1248 | chr2 | 89617266 | 89618190 | 0.220042098 | 0.016 |
| Gtpbp6 | chr5 | 110106652 | 110106687 | 0.220571953 | 0.016 |
| Zfp605 | chr5 | 110112051 | 110112066 | 0.220571953 | 0.016 |
| Atp8b4 | chr2 | 126358767 | 126358951 | 0.22113836 | 0.016 |
| Olfr13 | chr6 | 43173987 | 43174920 | 0.221868691 | 0.016 |
| Olfr1242 | chr2 | 89493389 | 89494310 | 0.221978384 | 0.016 |

**Supplementary Table 4. (continued).**

| | | | | | |
|---|---|---|---|---|---|
| Kcnu1 | chr8 | 25849655 | 25849850 | 0.222082301 | 0.016 |
| Ash2l | chr8 | 25831272 | 25831348 | 0.222082301 | 0.016 |
| Emc7 | chr2 | 112466980 | 112467133 | 0.225035524 | 0.016 |
| Nell1 | chr7 | 50701155 | 50701251 | 0.225577989 | 0.016 |
| Gm2115 | chr7 | 84576908 | 84577088 | 0.225671199 | 0.016 |
| Olfr1257 | chr2 | 89880827 | 89881757 | 0.225869269 | 0.016 |
| Xirp2 | chr2 | 67476809 | 67476866 | 0.229764986 | 0.016 |
| Ablim1 | chr19 | 57197298 | 57197314 | 0.229785163 | 0.016 |
| Lemd2 | chr17 | 27193102 | 27193204 | 0.230027116 | 0.016 |
| Mertk | chr2 | 128750099 | 128750186 | 0.230620746 | 0.016 |
| Itm2c | chr1 | 85905213 | 85905402 | 0.231162576 | 0.016 |
| Smoc2 | chr17 | 14348703 | 14348778 | 0.231241707 | 0.016 |
| Morc3 | chr16 | 93832309 | 93832348 | 0.231382002 | 0.016 |
| Cfap221 | chr1 | 119953600 | 119953709 | 0.232158789 | 0.017 |
| Rasgrf2 | chr13 | 92022265 | 92022287 | 0.232656399 | 0.017 |
| Olfr1196 | chr2 | 88700382 | 88701327 | 0.233393074 | 0.017 |
| Vwa8 | chr14 | 79009163 | 79009279 | 0.234565605 | 0.017 |
| Obscn | chr11 | 59054161 | 59054425 | 0.234962089 | 0.017 |
| Cntnap2 | chr6 | 47021569 | 47021788 | 0.235058012 | 0.017 |
| Zfp445 | chr9 | 122857469 | 122857638 | 0.236417523 | 0.017 |
| 2310002L09Rik | chr4 | 73939578 | 73939598 | 0.237256372 | 0.017 |
| Nmnat3 | chr9 | 98410076 | 98410441 | 0.237272135 | 0.017 |
| Olfr295 | chr7 | 86585276 | 86586206 | 0.238291174 | 0.017 |
| Olfr1393 | chr11 | 49280149 | 49281085 | 0.238415593 | 0.017 |
| Mgat1 | chr11 | 49260691 | 49262035 | 0.238415593 | 0.017 |
| Cdh13 | chr8 | 118505603 | 118505715 | 0.23856867 | 0.017 |
| Olfr1260 | chr2 | 89977779 | 89978712 | 0.239289248 | 0.017 |
| Fam76b | chr9 | 13836876 | 13836972 | 0.239610964 | 0.017 |
| Hist1h2be | chr13 | 23585575 | 23585956 | 0.240718218 | 0.017 |
| Olfr328 | chr11 | 58551304 | 58552237 | 0.241136251 | 0.017 |
| Olfr329-ps | chr11 | 58542517 | 58543483 | 0.241136251 | 0.017 |
| Plch1 | chr3 | 63850948 | 63850991 | 0.241298121 | 0.018 |
| E130311K13Rik | chr3 | 63929256 | 63929370 | 0.241671418 | 0.018 |
| Olfr141 | chr2 | 86806049 | 86806997 | 0.241914673 | 0.018 |
| Ptgs1 | chr2 | 36250995 | 36251351 | 0.242940458 | 0.018 |
| Folh1 | chr7 | 86762916 | 86763042 | 0.246203005 | 0.018 |
| Gtdc1 | chr2 | 44788901 | 44789043 | 0.246597769 | 0.018 |
| Tmem26 | chr10 | 68737098 | 68737102 | 0.247146462 | 0.018 |
| Olfr310 | chr7 | 86268788 | 86269787 | 0.249078977 | 0.018 |

**Supplementary Table 4. (continued).**

| | | | | | |
|---|---|---|---|---|---|
| Zfp712 | chr13 | 67040343 | 67042232 | 0.249424338 | 0.018 |
| Ifnab | chr4 | 88690654 | 88691227 | 0.250085612 | 0.018 |
| Olfr715 | chr7 | 107128443 | 107129391 | 0.250210193 | 0.018 |
| Unc5c | chr3 | 141733837 | 141733941 | 0.250627722 | 0.018 |
| Cd8b1 | chr6 | 71336709 | 71336722 | 0.252395138 | 0.018 |
| Mmp16 | chr4 | 17934069 | 17934114 | 0.253070034 | 0.019 |
| Clca4b | chr3 | 144911109 | 144911525 | 0.25374931 | 0.019 |
| Gemin5 | chr11 | 58141587 | 58141809 | 0.255109331 | 0.019 |
| Olfr1287 | chr2 | 111449141 | 111450059 | 0.255157849 | 0.019 |
| Vmn2r66 | chr7 | 85006522 | 85007329 | 0.255475084 | 0.019 |
| Olfr441 | chr6 | 43115743 | 43116676 | 0.255477016 | 0.019 |
| Trpm3 | chr19 | 22766633 | 22766808 | 0.256436466 | 0.019 |
| 9930021J03Rik | chr19 | 29786248 | 29786449 | 0.257327586 | 0.019 |
| Ralgapa1 | chr12 | 55794994 | 55795044 | 0.257748879 | 0.019 |
| Chrna7 | chr7 | 63148582 | 63148692 | 0.259657185 | 0.019 |
| Frg1 | chr8 | 41399564 | 41399656 | 0.260440132 | 0.019 |
| Cyp4f40 | chr17 | 32667815 | 32667869 | 0.260749719 | 0.019 |
| Olfr341 | chr2 | 36479186 | 36480128 | 0.261111654 | 0.019 |
| Olfr994 | chr2 | 85429882 | 85430827 | 0.261697409 | 0.019 |
| Olfr995 | chr2 | 85438208 | 85439156 | 0.261697409 | 0.019 |
| Tgm3 | chr2 | 130038362 | 130038608 | 0.261964124 | 0.019 |
| Olfr1298 | chr2 | 111645056 | 111645995 | 0.262106994 | 0.019 |
| Utp23 | chr15 | 51882150 | 51882169 | 0.263127675 | 0.020 |
| Nrxn3 | chr12 | 90204510 | 90204818 | 0.265891236 | 0.020 |
| Deptor | chr15 | 55252006 | 55252135 | 0.268175557 | 0.020 |
| Fabp1 | chr6 | 71199926 | 71199993 | 0.26851848 | 0.020 |
| Il1b | chr2 | 129365028 | 129365238 | 0.268998307 | 0.020 |
| Stk39 | chr2 | 68307040 | 68307108 | 0.270734969 | 0.020 |
| Olfr303 | chr7 | 86394536 | 86395496 | 0.271168582 | 0.020 |
| Olfr304 | chr7 | 86385656 | 86386658 | 0.271168582 | 0.020 |
| Armc9 | chr1 | 86202489 | 86202564 | 0.271385455 | 0.020 |
| Polr1b | chr2 | 129112641 | 129112813 | 0.271783534 | 0.020 |
| Chchd5 | chr2 | 129130177 | 129130318 | 0.271783534 | 0.020 |
| Serpinb3b | chr1 | 107154368 | 107154773 | 0.272400887 | 0.021 |
| Olfr859 | chr9 | 19808319 | 19809249 | 0.272563718 | 0.021 |
| Tasp1 | chr2 | 140048241 | 140048309 | 0.272664122 | 0.021 |
| Lsm8 | chr6 | 18850917 | 18851001 | 0.273886663 | 0.021 |
| Kctd18 | chr1 | 57967496 | 57967689 | 0.273936939 | 0.021 |
| Tmem217 | chr17 | 29526184 | 29526754 | 0.274123358 | 0.021 |

**Supplementary Table 4. (continued).**

| | | | | | |
|---|---|---|---|---|---|
| Igkv6-14 | chr6 | 70435427 | 70435476 | 0.274757225 | 0.021 |
| C1rb | chr6 | 124574256 | 124574403 | 0.275027668 | 0.021 |
| Olfr291 | chr7 | 84856364 | 84857318 | 0.277718164 | 0.021 |
| Asap1 | chr15 | 64208496 | 64208556 | 0.278823866 | 0.021 |
| Ckap2l | chr2 | 129269037 | 129269263 | 0.279679573 | 0.021 |
| Olfr1241 | chr2 | 89482188 | 89483133 | 0.280262703 | 0.021 |
| F830045P16Rik | chr2 | 129472634 | 129472949 | 0.280439906 | 0.021 |
| Rars | chr11 | 35828624 | 35828813 | 0.28078565 | 0.021 |
| Wwc1 | chr11 | 35839177 | 35839244 | 0.28078565 | 0.021 |
| Rgs7 | chr1 | 175059743 | 175059794 | 0.281062888 | 0.021 |
| Rrp1b | chr17 | 32060347 | 32060541 | 0.281107192 | 0.021 |
| Vmn2r-ps5 | chr3 | 64339564 | 64340547 | 0.28122093 | 0.021 |
| Eif3h | chr15 | 51865325 | 51865457 | 0.281841326 | 0.021 |
| Defb45 | chr2 | 152596338 | 152596393 | 0.282774109 | 0.022 |
| Defb36 | chr2 | 152604487 | 152604545 | 0.282774109 | 0.022 |
| Limk1 | chr5 | 134661793 | 134661858 | 0.283253306 | 0.022 |
| Fry | chr5 | 150405249 | 150405421 | 0.283338392 | 0.022 |
| Trim58 | chr11 | 58645895 | 58646126 | 0.283464311 | 0.022 |
| Gm13088 | chr4 | 143656399 | 143656647 | 0.285005436 | 0.022 |
| Zfp738 | chr13 | 67673561 | 67673688 | 0.286399092 | 0.022 |
| Vmn2r68 | chr7 | 85221517 | 85222437 | 0.287296658 | 0.022 |
| Sult6b2 | chr6 | 142797841 | 142797968 | 0.287611048 | 0.022 |
| Rnf17 | chr14 | 56459965 | 56460124 | 0.287752499 | 0.022 |
| Phlpp1 | chr1 | 106286990 | 106287116 | 0.287937859 | 0.022 |
| Chodl | chr16 | 78931368 | 78931447 | 0.289487657 | 0.022 |
| Klra5 | chr6 | 129911349 | 129911461 | 0.289943028 | 0.022 |
| Olfr301 | chr7 | 86412363 | 86413299 | 0.291013296 | 0.023 |
| Vmn1r180 | chr7 | 23952413 | 23953331 | 0.291484829 | 0.023 |
| Bfsp2 | chr9 | 103453061 | 103453144 | 0.291888247 | 0.023 |
| Try5 | chr6 | 41312186 | 41312316 | 0.292095583 | 0.023 |
| Tnfrsf8 | chr4 | 145288425 | 145288516 | 0.292189816 | 0.023 |
| Mfsd14b | chr13 | 65112503 | 65112617 | 0.292991748 | 0.023 |
| Carmil1 | chr13 | 24276456 | 24276554 | 0.293320022 | 0.023 |
| Sfta2 | chr17 | 35627993 | 35628163 | 0.293389709 | 0.023 |
| Gm9573 | chr17 | 35618621 | 35623231 | 0.293389709 | 0.023 |
| Mucl3 | chr17 | 35637355 | 35638623 | 0.293389709 | 0.023 |
| Trim30a | chr7 | 104429120 | 104429351 | 0.293410186 | 0.023 |
| Ctsj | chr13 | 61002402 | 61002565 | 0.293744092 | 0.023 |
| Rufy3 | chr5 | 88565148 | 88565476 | 0.294445665 | 0.023 |

**Supplementary Table 4. (continued).**

| | | | | | |
|---|---|---|---|---|---|
| Nox3 | chr17 | 3635264 | 3635391 | 0.294972816 | 0.023 |
| Nme6 | chr9 | 109839607 | 109839710 | 0.295676251 | 0.023 |
| Camp | chr9 | 109847434 | 109847581 | 0.295676251 | 0.023 |
| Zc3h8 | chr2 | 128935311 | 128935546 | 0.295969151 | 0.023 |
| Tnfrsf1b | chr4 | 145224251 | 145224484 | 0.296657179 | 0.023 |
| Eif3c | chr7 | 126556282 | 126556507 | 0.296912247 | 0.023 |
| Vmn2r56 | chr7 | 12693997 | 12694902 | 0.297318406 | 0.023 |
| Vmn2r55 | chr7 | 12684706 | 12684991 | 0.297318406 | 0.023 |
| Sprr2a3 | chr3 | 92288587 | 92288839 | 0.297684491 | 0.023 |
| Pacs2 | chr12 | 113062446 | 113062601 | 0.297800786 | 0.023 |
| Grm5 | chr7 | 87602543 | 87603201 | 0.298111149 | 0.023 |
| Olfr714 | chr7 | 107073829 | 107074783 | 0.298567356 | 0.023 |
| Cyp4f37 | chr17 | 32629034 | 32629156 | 0.300783498 | 0.024 |
| Mfn2 | chr4 | 147890113 | 147890276 | 0.300846993 | 0.024 |
| Ywhab | chr2 | 164014005 | 164014129 | 0.30139343 | 0.024 |
| Pabpc1l | chr2 | 164027474 | 164027668 | 0.30139343 | 0.024 |
| Olfr437 | chr6 | 43167059 | 43167992 | 0.301641657 | 0.024 |
| Olfr237-ps1 | chr6 | 43153306 | 43154239 | 0.301641657 | 0.024 |
| Olfr650-ps1 | chr7 | 104206028 | 104207009 | 0.302001895 | 0.024 |
| Cbfa2t2 | chr2 | 154500399 | 154500543 | 0.302740352 | 0.024 |
| Mtrr | chr13 | 68577657 | 68577705 | 0.303337121 | 0.024 |
| Olfr30 | chr11 | 58454999 | 58455947 | 0.303788814 | 0.024 |
| Gm12253 | chr11 | 58440956 | 58441056 | 0.303788814 | 0.024 |
| Lypd9 | chr11 | 58446207 | 58446401 | 0.303788814 | 0.024 |
| Plekhf1 | chr7 | 38221302 | 38222142 | 0.303927517 | 0.024 |
| H2-M2 | chr17 | 37482482 | 37482758 | 0.304934776 | 0.024 |
| Olfr110 | chr17 | 37498652 | 37499606 | 0.304934776 | 0.024 |
| Glo1 | chr17 | 30600038 | 30600179 | 0.305037684 | 0.024 |
| Lama3 | chr18 | 12557670 | 12557802 | 0.305702132 | 0.024 |
| Tenm4 | chr7 | 96694769 | 96695025 | 0.307072404 | 0.024 |
| Olfr1130 | chr2 | 87607389 | 87608334 | 0.308057742 | 0.024 |
| Olfr1094 | chr2 | 86828753 | 86829746 | 0.308490566 | 0.024 |
| Vmn2r75 | chr7 | 86163164 | 86163288 | 0.308892415 | 0.024 |
| Ipo9 | chr1 | 135402232 | 135402372 | 0.309362946 | 0.024 |
| Fpr1 | chr17 | 17876630 | 17877725 | 0.310046031 | 0.025 |
| Vmn2r25 | chr6 | 123851797 | 123852095 | 0.310344828 | 0.025 |
| Agbl4 | chr4 | 111617119 | 111617282 | 0.310475282 | 0.025 |
| Armc8 | chr9 | 99533097 | 99533190 | 0.310493314 | 0.025 |
| Slc5a12 | chr2 | 110624135 | 110624248 | 0.3107295 | 0.025 |

**Supplementary Table 4. (continued).**

| | | | | | |
|---|---|---|---|---|---|
| Dck | chr5 | 88772629 | 88772823 | 0.31076166 | 0.025 |
| Gm5724 | chr6 | 141725307 | 141725473 | 0.310810753 | 0.025 |
| 2900092C05Rik | chr7 | 12556012 | 12556053 | 0.311063258 | 0.025 |
| Tyr | chr7 | 87492645 | 87492890 | 0.311151532 | 0.025 |
| Stk24 | chr14 | 121337421 | 121337652 | 0.311305509 | 0.025 |
| Vmn2r53 | chr7 | 12584702 | 12584826 | 0.311498682 | 0.025 |
| Pde4b | chr4 | 102194835 | 102195074 | 0.311511537 | 0.025 |
| Nox4 | chr7 | 87338956 | 87339017 | 0.312087024 | 0.025 |
| Skint5 | chr4 | 113524095 | 113524158 | 0.312215241 | 0.025 |
| Iapp | chr6 | 142298848 | 142298931 | 0.312546383 | 0.025 |
| Uhrf1bp1 | chr17 | 27879223 | 27879413 | 0.31307588 | 0.025 |
| Vwa5b2 | chr16 | 20591357 | 20591390 | 0.313085645 | 0.025 |
| Olfr993 | chr2 | 85413932 | 85414877 | 0.313241026 | 0.025 |
| Map3k7 | chr4 | 31964262 | 31964382 | 0.31342103 | 0.025 |
| Tc2n | chr12 | 101688959 | 101689060 | 0.315090161 | 0.025 |
| Kcnk12 | chr17 | 87797063 | 87797454 | 0.315556243 | 0.025 |
| Ddx10 | chr9 | 53238047 | 53238178 | 0.316009207 | 0.025 |
| Olfr1057 | chr2 | 86374462 | 86375410 | 0.316250637 | 0.025 |
| Olfr1058 | chr2 | 86385465 | 86386416 | 0.316250637 | 0.025 |
| Tmc2 | chr2 | 130225794 | 130225876 | 0.316642433 | 0.025 |
| Ttl | chr2 | 129092940 | 129093055 | 0.316660782 | 0.025 |
| Enpp1 | chr10 | 24711805 | 24711991 | 0.316768594 | 0.025 |
| Erich2 | chr2 | 70523043 | 70523092 | 0.316926942 | 0.025 |
| Ss18 | chr18 | 14682666 | 14682735 | 0.317582725 | 0.025 |
| Slco1a6 | chr6 | 142086384 | 142086604 | 0.318163995 | 0.025 |
| Olfr300-ps1 | chr7 | 86442982 | 86443939 | 0.318363499 | 0.025 |
| Abt1 | chr13 | 23423585 | 23423832 | 0.318658189 | 0.025 |
| Impact | chr18 | 12975978 | 12976031 | 0.318998792 | 0.025 |
| Them5 | chr3 | 94344408 | 94344547 | 0.319679191 | 0.026 |
| Tuft1 | chr3 | 94658724 | 94658784 | 0.320011247 | 0.026 |
| Slc43a1 | chr2 | 84862882 | 84863029 | 0.320352223 | 0.026 |
| Wiz | chr17 | 32361460 | 32361964 | 0.32182241 | 0.026 |
| Serpinb3d | chr1 | 107079217 | 107079373 | 0.322047953 | 0.026 |
| Dnm3 | chr1 | 162320951 | 162321112 | 0.322062118 | 0.026 |
| Cmas | chr6 | 142764553 | 142764709 | 0.322282368 | 0.026 |
| Hikeshi | chr7 | 89923563 | 89923715 | 0.32359506 | 0.026 |
| Csn2 | chr5 | 87697119 | 87697140 | 0.326210506 | 0.026 |
| Ces1b | chr8 | 93079864 | 93079916 | 0.326809749 | 0.026 |
| Col9a1 | chr1 | 24196890 | 24196965 | 0.327408236 | 0.027 |

**Supplementary Table 4. (continued).**

| | | | | | |
|---|---|---|---|---|---|
| Adamts16 | chr13 | 70777388 | 70777484 | 0.328293356 | 0.027 |
| Vmn2r74 | chr7 | 85955917 | 85956145 | 0.328372476 | 0.027 |
| Klhl31 | chr9 | 77650003 | 77651175 | 0.328912842 | 0.027 |
| Zfp951 | chr5 | 104859666 | 104859696 | 0.329471084 | 0.027 |
| Pcdh9 | chr14 | 93560475 | 93560577 | 0.329648087 | 0.027 |
| Igkv4-63 | chr6 | 69377943 | 69378246 | 0.329702848 | 0.027 |
| Scgb1b24 | chr7 | 33743798 | 33743859 | 0.329788028 | 0.027 |
| Xrcc5 | chr1 | 72312407 | 72312591 | 0.330010046 | 0.027 |
| Tmem169 | chr1 | 72300683 | 72301306 | 0.330010046 | 0.027 |
| Ctsq | chr13 | 61039028 | 61039151 | 0.330180975 | 0.027 |
| Ift88 | chr14 | 57472974 | 57473074 | 0.330710257 | 0.027 |
| Epha1 | chr6 | 42372174 | 42372242 | 0.330731356 | 0.027 |
| Olfr1231 | chr2 | 89302648 | 89303590 | 0.33276273 | 0.027 |
| Olfr1230 | chr2 | 89296350 | 89297268 | 0.33276273 | 0.027 |
| Klra10 | chr6 | 130269209 | 130269345 | 0.332826667 | 0.027 |
| Stau2 | chr1 | 16440322 | 16440458 | 0.333481235 | 0.027 |
| Phactr3 | chr2 | 178119199 | 178119302 | 0.333791326 | 0.027 |
| Sult3a2 | chr10 | 33777169 | 33777264 | 0.334672759 | 0.028 |
| Cmah | chr13 | 24468546 | 24468622 | 0.335188256 | 0.028 |
| Fhit | chr14 | 9573412 | 9573511 | 0.335568469 | 0.028 |
| Olfr472 | chr7 | 107902718 | 107903651 | 0.335720975 | 0.028 |
| Grb14 | chr2 | 64938368 | 64938490 | 0.33598525 | 0.028 |
| Vmn2r67 | chr7 | 85155705 | 85155902 | 0.336314712 | 0.028 |
| Nsun7 | chr5 | 66294788 | 66294912 | 0.336695679 | 0.028 |
| Apbb2 | chr5 | 66302541 | 66302709 | 0.336695679 | 0.028 |
| Trps1 | chr15 | 50846175 | 50846940 | 0.336964659 | 0.028 |
| Cntnap5c | chr17 | 58055571 | 58055712 | 0.33731607 | 0.028 |
| Layn | chr9 | 51062182 | 51062266 | 0.338381201 | 0.028 |
| Acvr1 | chr2 | 58447538 | 58447673 | 0.339408686 | 0.028 |
| Trappc3l | chr10 | 34098763 | 34098949 | 0.339498563 | 0.028 |
| Calhm5 | chr10 | 34092125 | 34092515 | 0.339498563 | 0.028 |
| Dtnbp1 | chr13 | 44970045 | 44970178 | 0.340120783 | 0.028 |
| Tacstd2 | chr6 | 67534752 | 67535706 | 0.341151451 | 0.028 |
| Igkv4-51 | chr6 | 69681405 | 69681714 | 0.341214913 | 0.028 |
| Lrp1b | chr2 | 40725381 | 40725514 | 0.341312434 | 0.028 |
| Zfp128 | chr7 | 12881294 | 12881360 | 0.342278927 | 0.028 |
| Vmn2r73 | chr7 | 85875741 | 85875938 | 0.343050567 | 0.029 |
| Poc1a | chr9 | 106286779 | 106286887 | 0.343573696 | 0.029 |
| Smyd1 | chr6 | 71219257 | 71219421 | 0.343898765 | 0.029 |

**Supplementary Table 4. (continued).**

| | | | | | |
|---|---|---|---|---|---|
| Snapc3 | chr4 | 83453040 | 83453188 | 0.344179898 | 0.029 |
| Psip1 | chr4 | 83468582 | 83468676 | 0.344179898 | 0.029 |
| Mrps18c | chr5 | 100798769 | 100798860 | 0.344304837 | 0.029 |
| Helq | chr5 | 100785281 | 100785427 | 0.344304837 | 0.029 |
| Cog2 | chr8 | 124542888 | 124542950 | 0.344367938 | 0.029 |
| Fcgr4 | chr1 | 171025543 | 171025798 | 0.344594517 | 0.029 |
| Snx13 | chr12 | 35124429 | 35124540 | 0.344685989 | 0.029 |
| Vmn2r23 | chr6 | 123712365 | 123712419 | 0.34508936 | 0.029 |
| Gm10130 | chr2 | 150363002 | 150363071 | 0.345656147 | 0.029 |
| 3300002I08Rik | chr2 | 150362587 | 150362644 | 0.345656147 | 0.029 |
| Riox2 | chr16 | 59486532 | 59486635 | 0.345849005 | 0.029 |
| Crybg3 | chr16 | 59495609 | 59495768 | 0.345849005 | 0.029 |
| Irgm1 | chr11 | 48865800 | 48867003 | 0.345914512 | 0.029 |
| Zfp110 | chr7 | 12837088 | 12837184 | 0.346006258 | 0.029 |
| Gm13084 | chr4 | 143810736 | 143810857 | 0.346234351 | 0.029 |
| Pramel6 | chr2 | 87510200 | 87510783 | 0.346428333 | 0.029 |
| Olfr152 | chr2 | 87782535 | 87783486 | 0.346525983 | 0.029 |
| Asxl2 | chr12 | 3484444 | 3484608 | 0.347373293 | 0.029 |
| Prkacb | chr3 | 146747972 | 146748047 | 0.348524733 | 0.029 |
| Usp8 | chr2 | 126733959 | 126734104 | 0.348562955 | 0.029 |
| Usp50 | chr2 | 126733165 | 126733213 | 0.348562955 | 0.029 |
| Mptx1 | chr1 | 174332193 | 174332789 | 0.34856788 | 0.029 |
| Extl2 | chr3 | 116010688 | 116010696 | 0.349280096 | 0.029 |
| Slc30a7 | chr3 | 116006833 | 116006935 | 0.349280096 | 0.029 |
| Iqcb1 | chr16 | 36835515 | 36835645 | 0.34969998 | 0.029 |
| Muc13 | chr16 | 33807867 | 33808034 | 0.349945442 | 0.029 |
| Vmn2r72 | chr7 | 85737783 | 85738703 | 0.350477655 | 0.029 |
| Unc13b | chr4 | 43161442 | 43161478 | 0.350578871 | 0.029 |
| Rwdd1 | chr10 | 34019365 | 34019438 | 0.352837097 | 0.030 |
| Ipo7 | chr7 | 110018424 | 110018508 | 0.353300736 | 0.030 |
| Slco1a4 | chr6 | 141845408 | 141845468 | 0.35363512 | 0.030 |
| Cyp4f16 | chr17 | 32546368 | 32546435 | 0.35499523 | 0.030 |
| Ighg2c | chr12 | 113285324 | 113285408 | 0.355192833 | 0.030 |
| Ighe | chr12 | 113271709 | 113272030 | 0.355192833 | 0.030 |
| Gnl1 | chr17 | 35987724 | 35987887 | 0.355223615 | 0.030 |
| Aff4 | chr11 | 53380585 | 53380672 | 0.355346371 | 0.030 |
| Rbpms | chr8 | 33834303 | 33834454 | 0.355355739 | 0.030 |
| Polr2b | chr5 | 77332186 | 77332298 | 0.355493603 | 0.030 |
| Ctsm | chr13 | 61538383 | 61538546 | 0.355714795 | 0.030 |

**Supplementary Table 4. (continued).**

| | | | | | |
|---|---|---|---|---|---|
| Gm49352 | chr13 | 61538905 | 61539076 | 0.355714795 | 0.030 |
| Serpinb3c | chr1 | 107276852 | 107277014 | 0.356023287 | 0.030 |
| Magi2 | chr5 | 19227288 | 19227589 | 0.356113798 | 0.030 |
| Ddah1 | chr3 | 145845710 | 145845804 | 0.356234285 | 0.030 |
| Scn9a | chr2 | 66515314 | 66515432 | 0.357638778 | 0.030 |
| Ifi27l2b | chr12 | 103451074 | 103451352 | 0.357730857 | 0.030 |
| Parp9 | chr16 | 35947500 | 35948354 | 0.357765189 | 0.030 |
| Dtx3l | chr16 | 35938693 | 35938877 | 0.357765189 | 0.030 |
| Olfr305 | chr7 | 86363375 | 86364335 | 0.358016794 | 0.031 |
| Iqub | chr6 | 24450603 | 24450852 | 0.358432173 | 0.031 |
| Vmn2r69 | chr7 | 85412287 | 85412567 | 0.358435688 | 0.031 |
| Atp9b | chr18 | 80912796 | 80912947 | 0.358734413 | 0.031 |
| St8sia4 | chr1 | 95653512 | 95653770 | 0.358920031 | 0.031 |
| Ano3 | chr2 | 110713274 | 110713405 | 0.359603883 | 0.031 |
| Klra6 | chr6 | 130023637 | 130023727 | 0.360227009 | 0.031 |
| Olfr1008 | chr2 | 85689430 | 85690372 | 0.360244445 | 0.031 |
| Scgn | chr13 | 23966839 | 23966895 | 0.3607854 | 0.031 |
| Xkr6 | chr14 | 63660819 | 63660838 | 0.361189935 | 0.031 |
| Ddx24 | chr12 | 103424299 | 103424586 | 0.36128545 | 0.031 |
| Olfr1272 | chr2 | 90281646 | 90282573 | 0.362155478 | 0.031 |
| Abca13 | chr11 | 9451338 | 9451535 | 0.362210613 | 0.031 |
| Col14a1 | chr15 | 55408815 | 55408948 | 0.362233893 | 0.031 |
| Kyat3 | chr3 | 142744070 | 142744133 | 0.362356821 | 0.031 |
| Umodl1 | chr17 | 30954747 | 30954826 | 0.362383992 | 0.031 |
| Relch | chr1 | 105713848 | 105713939 | 0.363198074 | 0.031 |
| Olfr1205 | chr2 | 88831118 | 88832042 | 0.363240418 | 0.031 |
| Mrgpra9 | chr7 | 47252784 | 47252795 | 0.363457256 | 0.031 |
| Olfr322 | chr11 | 58665560 | 58666544 | 0.363548831 | 0.031 |
| Reep1 | chr6 | 71795146 | 71795324 | 0.363697599 | 0.031 |
| Ighd | chr12 | 113407534 | 113407543 | 0.363987517 | 0.031 |
| Il23r | chr6 | 67491768 | 67491780 | 0.364664872 | 0.031 |
| Spata3 | chr1 | 86026413 | 86026519 | 0.365205345 | 0.031 |
| Zfp459 | chr13 | 67408158 | 67408742 | 0.365646469 | 0.032 |
| Arhgap39 | chr15 | 76751527 | 76751947 | 0.365677359 | 0.032 |
| Tas2r122 | chr6 | 132710998 | 132711928 | 0.36660373 | 0.032 |
| Hist1h1c | chr13 | 23738848 | 23739487 | 0.367294976 | 0.032 |
| Stk38l | chr6 | 146773317 | 146773413 | 0.367299987 | 0.032 |
| Sp100 | chr1 | 85701116 | 85701139 | 0.368469164 | 0.032 |
| A630001G21Rik | chr1 | 85718192 | 85718368 | 0.368469164 | 0.032 |

**Supplementary Table 4. (continued).**

| | | | | | |
|---|---|---|---|---|---|
| Itprid2 | chr2 | 79651387 | 79651460 | 0.368470508 | 0.032 |
| Fbn1 | chr2 | 125335364 | 125335487 | 0.36888455 | 0.032 |
| Aard | chr15 | 52044798 | 52044954 | 0.368966099 | 0.032 |
| Hmgcll1 | chr9 | 76133754 | 76133865 | 0.369684376 | 0.032 |
| Olfr1115 | chr2 | 87251938 | 87252919 | 0.369883404 | 0.032 |
| Olfr1116 | chr2 | 87268845 | 87269769 | 0.369883404 | 0.032 |
| Hist1h4j | chr13 | 21735095 | 21735407 | 0.371528813 | 0.032 |
| Igsf21 | chr4 | 140027004 | 140027078 | 0.372177821 | 0.033 |
| Il21r | chr7 | 125622881 | 125622930 | 0.372495121 | 0.033 |
| Dnah9 | chr11 | 66005658 | 66005832 | 0.37257422 | 0.033 |
| Kifap3 | chr1 | 163815824 | 163815949 | 0.372920333 | 0.033 |
| Slc17a2 | chr13 | 23819017 | 23819136 | 0.372931836 | 0.033 |
| Ncr1 | chr7 | 4338122 | 4338366 | 0.373452061 | 0.033 |
| Sult2a4 | chr7 | 13984847 | 13984974 | 0.373564273 | 0.033 |
| Tas2r121 | chr6 | 132700089 | 132701007 | 0.373755198 | 0.033 |
| Oog4 | chr4 | 143450252 | 143450282 | 0.373974638 | 0.033 |
| Slco1a5 | chr6 | 142237521 | 142237586 | 0.375001897 | 0.033 |
| Btbd9 | chr17 | 30517103 | 30517323 | 0.375112564 | 0.033 |
| Atf7ip2 | chr16 | 10209133 | 10209268 | 0.37525355 | 0.033 |
| Rbfox1 | chr16 | 7330361 | 7330442 | 0.376172664 | 0.033 |
| Itgal | chr7 | 127314058 | 127314219 | 0.376297827 | 0.033 |
| Pip5k1a | chr3 | 95078101 | 95078182 | 0.37796899 | 0.033 |
| Kdm3a | chr6 | 71632055 | 71632241 | 0.378025273 | 0.033 |
| Polr1d | chr5 | 147078534 | 147078910 | 0.378537459 | 0.034 |
| Pyroxd1 | chr6 | 142356482 | 142356583 | 0.3792195 | 0.034 |
| Recql | chr6 | 142363546 | 142363676 | 0.3792195 | 0.034 |
| Bsph2 | chr7 | 13561331 | 13561364 | 0.379985246 | 0.034 |
| Cyp4f17 | chr17 | 32528046 | 32528111 | 0.380233853 | 0.034 |
| Olfr293 | chr7 | 86663663 | 86664674 | 0.380606148 | 0.034 |
| Hnmt | chr2 | 24003592 | 24003954 | 0.380834084 | 0.034 |
| Slc30a8 | chr15 | 52335121 | 52335264 | 0.381737387 | 0.034 |
| Pgm1 | chr4 | 99961455 | 99961618 | 0.382266938 | 0.034 |
| Olfr916 | chr9 | 38657457 | 38658390 | 0.382465218 | 0.034 |
| Olfr917 | chr9 | 38664912 | 38665842 | 0.382465218 | 0.034 |
| Larp1 | chr11 | 58009192 | 58009547 | 0.38260048 | 0.034 |
| Pif1 | chr9 | 65591719 | 65591859 | 0.382826086 | 0.034 |
| 1700009N14Rik | chr4 | 39450795 | 39451446 | 0.383192495 | 0.034 |
| Dpyd | chr3 | 119195068 | 119195188 | 0.383512912 | 0.034 |
| Ehmt2 | chr17 | 34898973 | 34899253 | 0.385202347 | 0.035 |

**Supplementary Table 4. (continued).**

| | | | | | |
|---|---|---|---|---|---|
| C2 | chr17 | 34881571 | 34881796 | 0.385202347 | 0.035 |
| Gm20547 | chr17 | 34881571 | 34881796 | 0.385202347 | 0.035 |
| Zbtb12 | chr17 | 34895240 | 34896620 | 0.385202347 | 0.035 |
| Spata48 | chr11 | 11496400 | 11496508 | 0.385823496 | 0.035 |
| Slco1a1 | chr6 | 141924335 | 141924500 | 0.386441051 | 0.035 |
| Golt1b | chr6 | 142392329 | 142392421 | 0.386754583 | 0.035 |
| Cdc7 | chr5 | 106975484 | 106975660 | 0.387426338 | 0.035 |
| Tas2r139 | chr6 | 42140935 | 42141895 | 0.387924171 | 0.035 |
| Olfr646 | chr7 | 104106280 | 104107219 | 0.387973041 | 0.035 |
| Nsmce1 | chr7 | 125486337 | 125486484 | 0.387977337 | 0.035 |
| Cks1brt | chr8 | 85171519 | 85171810 | 0.388178275 | 0.035 |
| Hdgfl1 | chr13 | 26769236 | 26770088 | 0.389731793 | 0.035 |
| Galnt18 | chr7 | 111471936 | 111472083 | 0.389815999 | 0.035 |
| 9130019O22Rik | chr7 | 127386519 | 127386633 | 0.39009845 | 0.035 |
| E430018J23Rik | chr7 | 127393252 | 127393436 | 0.39009845 | 0.035 |
| Ptprn2 | chr12 | 116858842 | 116859191 | 0.390544935 | 0.036 |
| St3gal5 | chr6 | 72149018 | 72149177 | 0.390931765 | 0.036 |
| Lamp3 | chr16 | 19699612 | 19699741 | 0.391076194 | 0.036 |
| Fkbpl | chr17 | 34645259 | 34645322 | 0.39113371 | 0.036 |
| Atf6b | chr17 | 34648218 | 34648297 | 0.39113371 | 0.036 |
| Rnpep | chr1 | 135264052 | 135264195 | 0.392015163 | 0.036 |
| Zfp799 | chr17 | 32822073 | 32822200 | 0.392565168 | 0.036 |
| Timd4 | chr11 | 46842155 | 46842195 | 0.392857594 | 0.036 |
| Clvs2 | chr10 | 33513258 | 33513346 | 0.392907732 | 0.036 |
| Aadacl3 | chr4 | 144463558 | 144463729 | 0.393782016 | 0.036 |
| Scube2 | chr7 | 109809077 | 109809245 | 0.393858641 | 0.036 |
| Olfr1286 | chr2 | 111420031 | 111420949 | 0.393989403 | 0.036 |
| Alcam | chr16 | 52274146 | 52274282 | 0.394174164 | 0.036 |
| Serinc3 | chr2 | 163625440 | 163625579 | 0.394263966 | 0.036 |
| Tmem108 | chr9 | 103484660 | 103484783 | 0.394395624 | 0.036 |
| Syne1 | chr10 | 5185395 | 5185557 | 0.394446583 | 0.036 |
| Ypel4 | chr2 | 84737419 | 84737528 | 0.394646205 | 0.036 |
| Gm19426 | chr2 | 84742470 | 84742700 | 0.394646205 | 0.036 |
| Susd1 | chr4 | 59349825 | 59350012 | 0.394840748 | 0.036 |
| Olfr318 | chr11 | 58720125 | 58721046 | 0.394962167 | 0.036 |
| Olfr317 | chr11 | 58732179 | 58733163 | 0.394962167 | 0.036 |
| Ubxn4 | chr1 | 128252202 | 128252305 | 0.394979265 | 0.036 |
| Ganc | chr2 | 120452592 | 120452667 | 0.395374264 | 0.036 |
| Kcna3 | chr3 | 107036422 | 107038009 | 0.395401022 | 0.036 |

**Supplementary Table 4. (continued).**

| | | | | | |
|---|---|---|---|---|---|
| Pramel7 | chr2 | 87492134 | 87492418 | 0.39541583 | 0.036 |
| Glyctk | chr9 | 106157488 | 106157865 | 0.395424547 | 0.036 |
| Ighg1 | chr12 | 113328975 | 113329298 | 0.395755335 | 0.036 |
| Rims3 | chr4 | 120889354 | 120889494 | 0.397304889 | 0.037 |
| Defb8 | chr8 | 19445859 | 19445984 | 0.397576887 | 0.037 |
| Pkig | chr2 | 163721150 | 163721198 | 0.397644866 | 0.037 |
| Ada | chr2 | 163730011 | 163730076 | 0.397644866 | 0.037 |
| Cpa6 | chr1 | 10477024 | 10477129 | 0.397881101 | 0.037 |
| Atp8b5 | chr4 | 43353546 | 43353709 | 0.397894867 | 0.037 |
| Zfp874b | chr13 | 67481718 | 67481845 | 0.398642716 | 0.037 |
| Prkch | chr12 | 73649291 | 73649355 | 0.399026858 | 0.037 |
| Gm6619 | chr6 | 131491406 | 131491435 | 0.399366315 | 0.037 |
| Olfr307 | chr7 | 86335452 | 86336394 | 0.399894578 | 0.037 |
| Bcl2l10 | chr9 | 75351010 | 75351133 | 0.399945958 | 0.037 |
| Slc33a1 | chr3 | 63944565 | 63944781 | 0.400050624 | 0.037 |
| Fzd6 | chr15 | 39030814 | 39031832 | 0.402260007 | 0.038 |
| Slc17a4 | chr13 | 23898854 | 23898869 | 0.402664973 | 0.038 |
| Slc17a1 | chr13 | 23892534 | 23892669 | 0.402664973 | 0.038 |
| Csta3 | chr16 | 36217662 | 36217788 | 0.402900987 | 0.038 |
| Lrrc55 | chr2 | 85191909 | 85192145 | 0.402919579 | 0.038 |
| Cep76 | chr18 | 67634754 | 67634940 | 0.403085803 | 0.038 |
| Psmg2 | chr18 | 67641678 | 67641735 | 0.403085803 | 0.038 |
| Slc26a8 | chr17 | 28638167 | 28638692 | 0.40317688 | 0.038 |
| Srpk1 | chr17 | 28622399 | 28622412 | 0.40317688 | 0.038 |
| Skint2 | chr4 | 112652057 | 112652224 | 0.40327892 | 0.038 |
| Hnf4a | chr2 | 163551576 | 163551751 | 0.403477312 | 0.038 |
| Arhgef12 | chr9 | 43040539 | 43040596 | 0.403860526 | 0.038 |
| Adam5 | chr8 | 24742097 | 24742151 | 0.404592714 | 0.038 |
| Pglyrp2 | chr17 | 32420799 | 32420863 | 0.404861399 | 0.038 |
| Vps13d | chr4 | 145181115 | 145181193 | 0.404969283 | 0.038 |
| Cfap299 | chr5 | 98498288 | 98498379 | 0.405621825 | 0.038 |
| Zfp874a | chr13 | 67443220 | 67443334 | 0.406210748 | 0.038 |
| Abi3bp | chr16 | 56686959 | 56687121 | 0.406534256 | 0.038 |
| Tfg | chr16 | 56694399 | 56694489 | 0.406534256 | 0.038 |
| Ndufa5 | chr6 | 24527272 | 24527317 | 0.40664176 | 0.038 |
| Vinac1 | chr2 | 129028068 | 129028186 | 0.406761101 | 0.038 |
| Olfr1132 | chr2 | 87634818 | 87635745 | 0.406893896 | 0.038 |
| Olfr1133 | chr2 | 87645179 | 87646121 | 0.406893896 | 0.038 |
| Gm5431 | chr11 | 48894334 | 48895546 | 0.407357466 | 0.039 |

**Supplementary Table 4. (continued).**

| | | | | | |
|---|---|---|---|---|---|
| Naa16 | chr14 | 79335733 | 79335931 | 0.407662939 | 0.039 |
| Nsun6 | chr2 | 15013139 | 15013284 | 0.408955656 | 0.039 |
| Hjv | chr3 | 96528063 | 96528690 | 0.409224459 | 0.039 |
| Tmem132a | chr19 | 10863954 | 10864104 | 0.409624654 | 0.039 |
| Tmem109 | chr19 | 10872586 | 10872689 | 0.409624654 | 0.039 |
| Olfr140 | chr2 | 90051413 | 90052322 | 0.410104498 | 0.039 |
| Stxbp5l | chr16 | 37115831 | 37116052 | 0.410140798 | 0.039 |
| C87436 | chr6 | 86457815 | 86457923 | 0.410337621 | 0.039 |
| Cdh6 | chr15 | 13053796 | 13053984 | 0.410455476 | 0.039 |
| Cep170 | chr1 | 176749925 | 176750052 | 0.410524993 | 0.039 |
| Mob1b | chr5 | 88749524 | 88749658 | 0.411157479 | 0.039 |
| Gm10272 | chr10 | 77706625 | 77706958 | 0.411364492 | 0.039 |
| Krtap12-1 | chr10 | 77720624 | 77721017 | 0.411364492 | 0.039 |
| Gm10024 | chr10 | 77711456 | 77711816 | 0.411364492 | 0.039 |
| Gm10142 | chr10 | 77715806 | 77716169 | 0.411364492 | 0.039 |
| Krit1 | chr5 | 3845482 | 3845507 | 0.411598746 | 0.040 |
| Lrrd1 | chr5 | 3849696 | 3851595 | 0.411598746 | 0.040 |
| Wrn | chr8 | 33329101 | 33329216 | 0.412580068 | 0.040 |
| Mis18bp1 | chr12 | 65161402 | 65161937 | 0.412683937 | 0.040 |
| Aspn | chr13 | 49566433 | 49566631 | 0.413244306 | 0.040 |
| Olfr1097 | chr2 | 86890225 | 86891173 | 0.413880628 | 0.040 |
| Gm13723 | chr2 | 86872773 | 86873721 | 0.413880628 | 0.040 |
| Hsd3b3 | chr3 | 98753310 | 98753455 | 0.413967552 | 0.040 |
| Epg5 | chr18 | 77959479 | 77959630 | 0.414521122 | 0.040 |
| Olfr1252 | chr2 | 89721164 | 89722109 | 0.415038472 | 0.040 |
| Islr | chr9 | 58156935 | 58158222 | 0.415243338 | 0.040 |
| Stra6 | chr9 | 58152444 | 58152600 | 0.415243338 | 0.040 |
| Gm14412 | chr2 | 177324304 | 177324307 | 0.415824753 | 0.040 |
| Ankrd31 | chr13 | 96821313 | 96821456 | 0.415964763 | 0.040 |
| Rhobtb3 | chr13 | 75939451 | 75939638 | 0.416160299 | 0.040 |
| Sptlc3 | chr2 | 139581513 | 139581619 | 0.416422549 | 0.040 |
| Dsel | chr1 | 111859179 | 111862803 | 0.416433455 | 0.040 |
| Kif1a | chr1 | 93082315 | 93082421 | 0.41684755 | 0.041 |
| Hist1h2aa | chr13 | 23934461 | 23934851 | 0.417046329 | 0.041 |
| Hist1h2ba | chr13 | 23933772 | 23934156 | 0.417046329 | 0.041 |
| Zfp458 | chr13 | 67269030 | 67269036 | 0.417482978 | 0.041 |
| Serpinb8 | chr1 | 107602799 | 107602917 | 0.417812025 | 0.041 |
| Adamts10 | chr17 | 33549479 | 33549609 | 0.418821659 | 0.041 |
| Myo1f | chr17 | 33555853 | 33555856 | 0.418821659 | 0.041 |

**Supplementary Table 4. (continued).**

| | | | | | |
|---|---|---|---|---|---|
| Lrpap1 | chr5 | 35105478 | 35105691 | 0.418880688 | 0.041 |
| Rbm47 | chr5 | 66024952 | 66025159 | 0.419161671 | 0.041 |
| Myh15 | chr16 | 49091074 | 49091178 | 0.419182653 | 0.041 |
| Olfr713 | chr7 | 107036135 | 107037110 | 0.419217364 | 0.041 |
| Slc5a9 | chr4 | 111893167 | 111893332 | 0.420069146 | 0.041 |
| Trbv21 | chr6 | 41202754 | 41203044 | 0.420348519 | 0.041 |
| Olfr1394 | chr11 | 49160015 | 49160954 | 0.420627663 | 0.041 |
| Olfr1395 | chr11 | 49148258 | 49149212 | 0.420627663 | 0.041 |
| Pbld1 | chr10 | 63075060 | 63075123 | 0.421794393 | 0.041 |
| Smim14 | chr5 | 65453185 | 65453328 | 0.422008628 | 0.041 |
| Ankub1 | chr3 | 57657436 | 57657566 | 0.423416909 | 0.042 |
| Commd2 | chr3 | 57650207 | 57650215 | 0.423416909 | 0.042 |
| Madd | chr2 | 91155545 | 91155648 | 0.423733421 | 0.042 |
| Il1a | chr2 | 129306463 | 129306692 | 0.423875545 | 0.042 |
| Nemf | chr12 | 69337818 | 69337907 | 0.424025731 | 0.042 |
| Il4ra | chr7 | 125571933 | 125571953 | 0.425129702 | 0.042 |
| Prkcz | chr4 | 155286779 | 155286877 | 0.426109767 | 0.042 |
| Brca2 | chr5 | 150548488 | 150548835 | 0.426166771 | 0.042 |
| Dtna | chr18 | 23595458 | 23595625 | 0.426773677 | 0.042 |
| Stoml3 | chr3 | 53500751 | 53500834 | 0.427138416 | 0.042 |
| Frem2 | chr3 | 53516531 | 53517041 | 0.427138416 | 0.042 |
| Fut10 | chr8 | 31201209 | 31201504 | 0.427663883 | 0.043 |
| Pdik1l | chr4 | 134284245 | 134284530 | 0.427707188 | 0.043 |
| Sit1 | chr4 | 43483537 | 43483583 | 0.427934 | 0.043 |
| Ralb | chr1 | 119471705 | 119471825 | 0.428225009 | 0.043 |
| Olfr1177-ps | chr2 | 88346262 | 88346291 | 0.428396306 | 0.043 |
| Olfr1176 | chr2 | 88339566 | 88340514 | 0.428396306 | 0.043 |
| Olfr1006 | chr2 | 85674210 | 85675149 | 0.429552179 | 0.043 |
| Raly | chr2 | 154859677 | 154859750 | 0.429799967 | 0.043 |
| Ighg2b | chr12 | 113306823 | 113307153 | 0.430298387 | 0.043 |
| Sag | chr1 | 87805311 | 87805377 | 0.430655292 | 0.043 |
| Rock1 | chr18 | 10150234 | 10150316 | 0.430669117 | 0.043 |
| D2hgdh | chr1 | 93829756 | 93829896 | 0.431174552 | 0.043 |
| S100a1 | chr3 | 90511978 | 90512136 | 0.431618 | 0.043 |
| Chtop | chr3 | 90499901 | 90500107 | 0.431618 | 0.043 |
| S100a13 | chr3 | 90515849 | 90516042 | 0.431618 | 0.043 |
| S100a8 | chr3 | 90669541 | 90669682 | 0.431922741 | 0.043 |
| Rad21 | chr15 | 51976098 | 51976198 | 0.43195269 | 0.043 |
| Vmn1r30 | chr6 | 58434936 | 58435845 | 0.432039798 | 0.043 |

**Supplementary Table 4. (continued).**

| | | | | | |
|---|---|---|---|---|---|
| Anapc1 | chr2 | 128666564 | 128666705 | 0.43256557 | 0.043 |
| Slc17a6 | chr7 | 51649151 | 51649187 | 0.432983182 | 0.044 |
| Olfm4 | chr14 | 80021118 | 80021930 | 0.433290782 | 0.044 |
| Vipr2 | chr12 | 116094720 | 116094818 | 0.433463358 | 0.044 |
| Zfp791 | chr8 | 85112209 | 85112270 | 0.433511955 | 0.044 |
| Pxmp4 | chr2 | 154603516 | 154603629 | 0.43357214 | 0.044 |
| Nfib | chr4 | 82310302 | 82310391 | 0.433781414 | 0.044 |
| Pld5 | chr1 | 176089870 | 176090039 | 0.434226813 | 0.044 |
| Usp39 | chr6 | 72339966 | 72340061 | 0.434262011 | 0.044 |
| 0610030E20Rik | chr6 | 72348996 | 72349119 | 0.434262011 | 0.044 |
| Retnlg | chr16 | 48874216 | 48874344 | 0.435429909 | 0.044 |
| Wdr19 | chr5 | 65202559 | 65202651 | 0.435754959 | 0.044 |
| Abcc9 | chr6 | 142590365 | 142590503 | 0.435884408 | 0.044 |
| 1700012P22Rik | chr4 | 144438487 | 144438604 | 0.436091267 | 0.044 |
| Gm12248 | chr11 | 58063810 | 58063976 | 0.436752483 | 0.044 |
| Col11a1 | chr3 | 114105389 | 114105446 | 0.437415578 | 0.044 |
| Tjp1 | chr7 | 65370809 | 65370836 | 0.437839908 | 0.044 |
| Ugt1a7c | chr1 | 88095120 | 88095978 | 0.437947726 | 0.044 |
| Ugt1a8 | chr1 | 88087866 | 88088721 | 0.437947726 | 0.044 |
| Pde4d | chr13 | 108860165 | 108860207 | 0.438219808 | 0.044 |
| Rab7 | chr6 | 88004992 | 88005211 | 0.438674196 | 0.044 |
| Atpaf1 | chr4 | 115811001 | 115811196 | 0.438716903 | 0.045 |
| Ldhb | chr6 | 142490587 | 142490604 | 0.438744463 | 0.045 |
| Gys2 | chr6 | 142472680 | 142472801 | 0.438744463 | 0.045 |
| C1qtnf4 | chr2 | 90889384 | 90890365 | 0.438795808 | 0.045 |
| Akap8 | chr17 | 32321084 | 32321103 | 0.439032385 | 0.045 |
| Akap8l | chr17 | 32336284 | 32336738 | 0.439032385 | 0.045 |
| Rbpms2 | chr9 | 65630828 | 65630897 | 0.439131001 | 0.045 |
| Olfr1232 | chr2 | 89325242 | 89326178 | 0.43925819 | 0.045 |
| Crocc2 | chr1 | 93181072 | 93181221 | 0.439645814 | 0.045 |
| Fntb | chr12 | 76862485 | 76862535 | 0.439679805 | 0.045 |
| Cntn3 | chr6 | 102269103 | 102269240 | 0.439769588 | 0.045 |
| Spx | chr6 | 142414020 | 142414078 | 0.439887607 | 0.045 |
| B3galt1 | chr2 | 68117942 | 68118923 | 0.440120869 | 0.045 |
| Ppox | chr1 | 171280673 | 171280760 | 0.440194634 | 0.045 |
| Ufc1 | chr1 | 171294633 | 171294756 | 0.440194634 | 0.045 |
| Usp21 | chr1 | 171284926 | 171285038 | 0.440194634 | 0.045 |
| Tmem67 | chr4 | 12073893 | 12073956 | 0.440308776 | 0.045 |
| Chfr | chr5 | 110136061 | 110136194 | 0.440650758 | 0.045 |

**Supplementary Table 4. (continued).**

| | | | | | |
|---|---|---|---|---|---|
| Zfp672 | chr11 | 58329417 | 58329771 | 0.441534134 | 0.045 |
| Cstdc6 | chr16 | 36321721 | 36321847 | 0.441612859 | 0.045 |
| Mbtps1 | chr8 | 119528923 | 119529068 | 0.441684401 | 0.045 |
| Sbf2 | chr7 | 110330609 | 110330711 | 0.441769242 | 0.045 |
| Slc28a1 | chr7 | 81124867 | 81125009 | 0.441873601 | 0.045 |
| Oc90 | chr15 | 65900590 | 65900647 | 0.442008217 | 0.045 |
| Cts8 | chr13 | 61250906 | 61251069 | 0.442747907 | 0.046 |
| Prpf6 | chr2 | 181631930 | 181632087 | 0.443011319 | 0.046 |
| Gpc5 | chr14 | 115788172 | 115788332 | 0.443071139 | 0.046 |
| Ugt2b36 | chr5 | 87066437 | 87066470 | 0.443279548 | 0.046 |
| Prr27 | chr5 | 87841257 | 87841281 | 0.443332815 | 0.046 |
| Csn1s2b | chr5 | 87824138 | 87824144 | 0.443332815 | 0.046 |
| Rasal3 | chr17 | 32391335 | 32391399 | 0.443469867 | 0.046 |
| Olfr173 | chr16 | 58796878 | 58797844 | 0.443755577 | 0.046 |
| Olfr988 | chr2 | 85352994 | 85353924 | 0.444361783 | 0.046 |
| Sephs2 | chr7 | 127272560 | 127273919 | 0.446248804 | 0.046 |
| Lrrc7 | chr3 | 158353264 | 158353467 | 0.447616049 | 0.047 |
| Olfr536 | chr7 | 140503533 | 140504457 | 0.447627118 | 0.047 |
| Fcgr3 | chr1 | 171058585 | 171058606 | 0.44795498 | 0.047 |
| Tram2 | chr1 | 21037396 | 21037460 | 0.448012735 | 0.047 |
| Aktip | chr8 | 91129599 | 91129805 | 0.448133224 | 0.047 |
| Rbl2 | chr8 | 91115040 | 91115249 | 0.448133224 | 0.047 |
| Bdh2 | chr3 | 135296819 | 135296933 | 0.448287427 | 0.047 |
| Smr2 | chr5 | 88088190 | 88088244 | 0.449103235 | 0.047 |
| Pramel1 | chr4 | 143397049 | 143397616 | 0.449373617 | 0.047 |
| Ppt1 | chr4 | 122845970 | 122846041 | 0.449503732 | 0.047 |
| Hs2st1 | chr3 | 144556981 | 144557085 | 0.449698947 | 0.047 |
| Klri1 | chr6 | 129703314 | 129703380 | 0.449813011 | 0.047 |
| Olfr366 | chr2 | 37219490 | 37220420 | 0.449901318 | 0.047 |
| Ago4 | chr4 | 126509950 | 126510135 | 0.450009348 | 0.047 |
| Esyt2 | chr12 | 116334200 | 116334256 | 0.450559871 | 0.047 |
| Slc38a6 | chr12 | 73352456 | 73352537 | 0.45086383 | 0.047 |
| Igtp | chr11 | 58206043 | 58206119 | 0.450903341 | 0.047 |
| Olfr1111 | chr2 | 87149720 | 87150659 | 0.451471324 | 0.047 |
| Chmp3 | chr6 | 71559013 | 71559033 | 0.45152406 | 0.047 |
| Gtf3c1 | chr7 | 125642446 | 125642639 | 0.451627712 | 0.047 |
| Klhl5 | chr5 | 65143238 | 65143375 | 0.451789586 | 0.047 |
| Olfr452 | chr6 | 42790040 | 42790994 | 0.451826166 | 0.048 |
| Olfr1330 | chr4 | 118893084 | 118894032 | 0.451943194 | 0.048 |

**Supplementary Table 4. (continued).**

| | | | | | |
|---|---|---|---|---|---|
| 4930438A08Rik | chr11 | 58286494 | 58286724 | 0.452147237 | 0.048 |
| Gm13757 | chr2 | 88446018 | 88446936 | 0.452457482 | 0.048 |
| Nme7 | chr1 | 164385699 | 164385801 | 0.452474553 | 0.048 |
| Trim66 | chr7 | 109472233 | 109472395 | 0.452499618 | 0.048 |
| Skint9 | chr4 | 112390911 | 112391016 | 0.452715924 | 0.048 |
| Akap13 | chr7 | 75602785 | 75602978 | 0.452736913 | 0.048 |
| Ccdc178 | chr18 | 22018980 | 22019126 | 0.454159331 | 0.048 |
| Btn1a1 | chr13 | 23458702 | 23459367 | 0.454450364 | 0.048 |
| Pank2 | chr2 | 131287473 | 131287597 | 0.454537408 | 0.048 |
| Gabrr2 | chr4 | 33085545 | 33085742 | 0.454670323 | 0.048 |
| Wdr95 | chr5 | 149595249 | 149595391 | 0.454705495 | 0.048 |
| Synpr | chr14 | 13285140 | 13285206 | 0.454720181 | 0.048 |
| Vmn1r223 | chr13 | 23249237 | 23250323 | 0.454761825 | 0.048 |
| Fam151a | chr4 | 106743196 | 106743356 | 0.454974055 | 0.048 |
| Acot11 | chr4 | 106749261 | 106749388 | 0.454974055 | 0.048 |
| Dpy19l2 | chr9 | 24691955 | 24691980 | 0.455014017 | 0.048 |
| Stk32b | chr5 | 37490460 | 37490550 | 0.455566844 | 0.048 |
| Thap4 | chr1 | 93705535 | 93705655 | 0.455627333 | 0.048 |
| Slc17a3 | chr13 | 23846210 | 23846444 | 0.455802001 | 0.048 |
| Sult2a8 | chr7 | 14413628 | 14413797 | 0.456085792 | 0.048 |
| Htr2b | chr1 | 86100100 | 86100232 | 0.45625254 | 0.048 |
| Psmd1 | chr1 | 86116574 | 86116689 | 0.45625254 | 0.048 |
| Olfr504 | chr7 | 108564836 | 108565793 | 0.456315495 | 0.048 |
| Myo5b | chr18 | 74569766 | 74569877 | 0.456740833 | 0.048 |
| Olfr1166 | chr2 | 88124032 | 88124983 | 0.457317923 | 0.048 |
| Nrg3 | chr14 | 38509329 | 38509356 | 0.457457646 | 0.048 |
| 4930470P17Rik | chr2 | 170579490 | 170579831 | 0.457807857 | 0.049 |
| Galnt17 | chr5 | 131111688 | 131111855 | 0.457958018 | 0.049 |
| Usp17lb | chr7 | 104842476 | 104842504 | 0.458128079 | 0.049 |
| 3110040N11Rik | chr7 | 81783042 | 81783138 | 0.458306175 | 0.049 |
| Ramac | chr7 | 81768368 | 81768558 | 0.458306175 | 0.049 |
| Cchcr1 | chr17 | 35530700 | 35530834 | 0.458778082 | 0.049 |
| Psors1c2 | chr17 | 35533239 | 35533294 | 0.458778082 | 0.049 |
| Olfr56 | chr11 | 49134289 | 49135141 | 0.458866446 | 0.049 |
| Tas2r123 | chr6 | 132847141 | 132848143 | 0.458958794 | 0.049 |
| Ptprd | chr4 | 76243644 | 76243786 | 0.459015796 | 0.049 |
| Rp1 | chr1 | 4206659 | 4206837 | 0.459522423 | 0.049 |
| Prss47 | chr13 | 65049214 | 65049492 | 0.459648964 | 0.049 |
| Kcnh1 | chr1 | 192434739 | 192434919 | 0.459870378 | 0.049 |

**Supplementary Table 4. (continued).**

| | | | | | |
|---|---|---|---|---|---|
| Slc2a2 | chr3 | 28726230 | 28726332 | 0.460056246 | 0.049 |
| Snap47 | chr11 | 59407663 | 59407810 | 0.460158827 | 0.049 |
| Ufl1 | chr4 | 25259214 | 25259337 | 0.460720951 | 0.049 |
| Slco6c1 | chr1 | 97104787 | 97104906 | 0.460747626 | 0.049 |
| Ifna5 | chr4 | 88835524 | 88836094 | 0.460963428 | 0.049 |
| Ifna4 | chr4 | 88841860 | 88842421 | 0.460963428 | 0.049 |
| Spi1 | chr2 | 91096948 | 91096993 | 0.46132249 | 0.049 |
| Prrc2a | chr17 | 35162201 | 35162379 | 0.461865109 | 0.049 |
| Aif1 | chr17 | 35172284 | 35172346 | 0.461865109 | 0.049 |
| Fam114a1 | chr5 | 64995814 | 64995902 | 0.462004551 | 0.049 |
| Zfp708 | chr13 | 67074085 | 67074181 | 0.462061157 | 0.049 |
| Olfr1245 | chr2 | 89574800 | 89575724 | 0.462537615 | 0.049 |
| Olfr1246 | chr2 | 89590168 | 89591113 | 0.462537615 | 0.049 |

* Genomic position is based on GRCm38 (mm10)

** the level of the subspecies musculus genomic ancestry in the window

*** Adjusted p values using Benjamini-Hochberg method
